# Quorum Sensing Regulates ‘swim-or-stick’ Lifestyle in the Phycosphere

**DOI:** 10.1101/2020.07.26.221937

**Authors:** Cong Fei, Michael A. Ochsenkühn, Ahmed A. Shibl, Ashley Isaac, Changhai Wang, Shady A. Amin

**Author notes:** **Corresponding author:** Dr. Shady. A. Amin, Marine Microbial Ecology Lab, Biology Program, New York University Abu Dhabi, Abu Dhabi, United Arab Emirates. (SAA), PO Box 129188, NYU Abu Dhabi, Abu Dhabi, UAE.

## Abstract

Motility and biofilm formation are processes regulated by quorum sensing (QS) in bacteria. Both functions are believed to play an important role in interactions between bacteria and phytoplankton. Here, we show that two bacterial symbionts from the microbial community associated with a ubiquitous diatom switch their motile lifestyle to attached cells while an opportunist bacterium from the same community is incapable of attachment, despite possessing the genetic machinery to do so. Further work indicated that the opportunist lacks QS signal synthases while the symbionts produce three QS signals, one of which is mainly responsible for regulating symbiont colonization of the diatom microenvironment. These findings suggest that QS regulates colonization of diatom surfaces and further work on these model systems will inform our understanding of particle aggregation and bacterial attachment to marine snow and how these processes influence the global carbon cycle.

**Summary:** Interactions between phytoplankton and bacteria play major roles in global biogeochemical cycles and oceanic nutrient fluxes. These interactions occur in the microenvironment surrounding phytoplankton cells, known as the phycosphere. Bacteria in the phycosphere use either chemotaxis or attachment to benefit from algal excretions. Both processes are regulated by quorum sensing (QS), a cell-cell signaling mechanism that uses small infochemicals to coordinate bacterial gene expression. However, the role of QS in regulating bacterial attachment in the phycosphere is not clear. Here, we isolated a *Sulfitobacter pseudonitzschiae* F5 and a *Phaeobacter* sp. F10 belonging to the marine *Roseobacter* group and an *Alteromonas macleodii* F12 belonging to Alteromonadaceae, from the microbial community of the ubiquitous diatom *Asterionellopsis glacialis.* We show that only the *Roseobacter* group isolates (diatom symbionts) can attach to diatom transparent exopolymeric particles. Despite all three bacteria possessing genes involved in motility, chemotaxis, and attachment, only *S. pseudonitzschiae* F5 and *Phaeobacter* sp. F10 possessed complete QS systems and could synthesize QS signals. Using UHPLC-MS/MS, we identified three QS molecules produced by both bacteria of which only 3-oxo-C_16:1_-HSL strongly inhibited bacterial motility and stimulated attachment in the phycosphere. These findings suggest that QS signals enable colonization of the phycosphere by algal symbionts.

## Introduction

Phytoplankton constitute the foundation of the marine food web as they are responsible for nearly half of primary production on Earth (Field et al., 1998). Through their ability to carry out photosynthesis, phytoplankton transform atmospheric carbon dioxide gas to organic matter (Simon et al., 2009) that is assimilated and remineralized by heterotrophic bacteria (Pomeroy, 1974; Burkhardt et al., 2014). In exchange, bacteria produce essential factors (e.g., vitamins) (Kazamia et al., 2012; Bertrand et al., 2015) to support the growth of phytoplankton (Amin et al., 2012). Cumulatively, phytoplankton-bacteria symbiosis is believed to play an important role in nutrient availability and major biogeochemical cycles (Buchan et al., 2014; Durham et al., 2019).

Several bacterial lineages have been consistently observed to co-occur with phytoplankton, such as members of the *Roseobacter* group, Gammaproteobacteria, and Flavobacteria (Wagner-Döbler and Biebl, 2006; Teeling et al., 2012). Particularly, members of the *Roseobacter* group (hereafter roseobacters) have been shown consistently to form symbiotic relationships with phytoplankton (Amin et al., 2012). For example, roseobacters are adept at acquiring and assimilating phytoplankton metabolites (Miller et al., 2004) in exchange for a variety of cofactors. *Ruegeria pomeroyi* has been shown to assimilate organic sulfur compounds from the diatom *Thalassiosira pseudonana* in exchange for production of cobalamin, which is required for diatom growth (Durham et al., 2015). *Sulfitobacter pseudonitzschiae* SA11 produces the hormone indole-3-acetic acid (IAA) to enhance cell division of the diatom *Pseudo-nitzschia multiseries*, which leads to an increase in carbon export by the diatom and the exchange of diatom-derived organosulfur compounds and bacterial ammonia (Amin et al., 2015). Other phytoplankton such as the coccolithophore, *Emiliania huxleyi*, display different morphologies in response to endogenous IAA (Labeeuw et al., 2016) and exhibit enhanced cell division by roseobacters-derived IAA (Segev et al., 2016; Bramucci et al., 2018). IAA is an endogenous hormone that regulates plant differentiation and is also produced and excreted by plant symbionts and some plant pathogens to interfere with plant root differentiation (Spaepen et al., 2007; Spaepen and Vanderleyden, 2011). Although eukaryotic phytoplankton like diatoms and coccolithophores are unicellular and do not undergo differentiation as defined in multicellular eukaryotes, IAA appears to have evolved the ability to manipulate the cell cycle in both groups of organisms, with related bacteria playing an important role in eukaryotic IAA perception in both marine and rhizobial environments (Seymour et al., 2017). A prerequisite for these symbiotic exchanges to occur is the intimate spatial proximity between the phytoplankton host and its symbionts.

Chemical exchanges between bacteria and phytoplankton occur in the microenvironment immediately adjacent to phytoplankton cells, known as the phycosphere (Bell and Mitchell, 1972; Seymour et al., 2017). Phytoplankton continuously exude organic matter, sometimes up to 50% of the cell’s total fixed carbon (Thornton, 2014). Consequently, the phycosphere is hypothesized to harbor a significantly higher amount of phytoplankton dissolved organic matter (DOM) relative to bulk seawater due to the exudation of DOM by phytoplankton cells, and due to the negligible effects of turbulence on the diffusion of exudates within the minute phycosphere (Seymour et al., 2017). This buildup of DOM ultimately leads to bacterial attraction and colonization of the phycosphere either via chemotaxis or random encounters with phytoplankton cells (Smriga et al., 2016). Once in the phycosphere, beneficial bacteria that produce metabolites essential to phytoplankton (Mayali et al., 2011) may gain an advantage by switching their free-living, planktonic lifestyle in bulk seawater to a surface-attached state on phytoplankton cells. On the other hand, opportunistic bacteria benefit from phytoplankton-derived DOM without providing apparent benefits to phytoplankton hosts (Mayali and Doucette, 2002). Compared to free-living cells, surface-associated cells have greater access to phytoplankton nutrients and gain protection against toxins, antibiotics, and other environmental stressors by forming a biofilm (Jefferson, 2004; Samo et al., 2018).

Successful colonization of the phycosphere can be enhanced by specific bacterial genetic traits, including chemotaxis, motility, and attachment to surfaces (Slightom and Buchan, 2009; Raina et al., 2019). Chemotaxis enables bacteria to sense changes in local concentrations of food and regulates motility accordingly. Motile bacteria generally exhibit strong chemotaxis to DOM released from phytoplankton (Miller and Belas, 2004; Miller et al., 2004; Stocker, 2012; Smriga et al., 2016), while motility and flagellar genes appear to be critical for attachment and biofilm development in many roseobacters (Miller and Belas, 2006; Bruhn et al., 2007). In the phycosphere, many bacteria may have a biphasic ‘swim-or-stick’ lifestyle that enables them to rapidly find food sources while minimizing energy expenditures once the food is located. During the motile phase, bacteria use chemotaxis to locate phytoplankton cells (Seymour et al., 2017). Once in the phycosphere, a ‘switch’ is turned on, causing a transition of the bacteria to a sessile lifestyle, whose phenotype includes loss of flagella and subsequent biofilm development (Geng and Belas, 2010). Despite our knowledge of bacterial behavior, the mechanisms that regulate bacterial motility and attachment in the phycosphere have not been extensively investigated.

Quorum sensing (QS) is one of the best-studied signaling mechanisms for bacterial cell-to-cell communication. Many bacteria have been shown to carry out QS by secreting small signaling molecules, known as autoinducers, to assess changes in bacterial populations and to coordinate gene expression among a whole population (Waters and Bassler, 2005). In Proteobacteria, the primary class of autoinducers is acyl-homoserine lactones (AHLs), which are synthesized by an autoinducer synthase (LuxI) and perceived by an autoinducer regulator (LuxR) (Case et al., 2008). Bacteria use AHLs to regulate functions that are beneficial to carry out collectively, such as virulence, motility, and biofilm formation (Hammer and Bassler, 2003; Daniels et al., 2004; Antunes et al., 2010). Indeed, several *Roseobacter*-group bacteria use AHLs to regulate motility, virulence, biofilm formation, and nutrient acquisition when associated with marine snow (Gram et al., 2002; Hmelo et al., 2011), red alga (Gardiner et al., 2015), and sponges (Zan et al., 2012). The addition of exogenous AHLs produced by bacterial epibionts to colonies of the cyanobacterium *Trichodesmium* led to increases in the activity of alkaline phosphatase activity and consequently phosphorus acquisition (Van Mooy et al., 2012). In contrast, the lack of complete QS systems often leads to the inability of bacteria to attach to hosts. For example, a *luxR*-type gene knock-out strain of the *Roseobacter*-group member *Nautella italica* R11 was unable to form biofilms or attach to the red alga *Delisea pulchra* (Gardiner et al., 2015). Despite these examples, there is little direct evidence showing how QS regulates bacteria-phytoplankton interactions or how QS influences inter- and intraspecies interactions and behavior between bacteria within microbial consortia in the phycosphere. In this study, we examine how QS influences bacterial behavior in the phycosphere of the ubiquitous diatom, *Asterionellopsis glacialis*, and whether QS provides an advantage to beneficial bacteria relative to other bacteria. Here we hypothesize that QS regulates motility and attachment of beneficial bacteria, which may enhance their access to phytoplankton nutrients over non-beneficial bacteria.

*A. glacialis* is a ubiquitous diatom that has been isolated from every major water body around the world (Korner, 1970; Kaczmarska et al., 2014) and has recently been shown to be one of several abundant and widely distributed groups of diatoms from the Tara Oceans expedition (Malviya et al., 2016). In addition, *A. glacialis* often forms blooms and dense patches worldwide (Karentz and Smayda, 1984; Franco et al., 2016) that are characterized by high DOM secretions (Abreu et al., 2003), making this diatom an ideal model system to examine interactions with bacteria. Here, we characterize AHL molecules produced by bacteria isolated from the phycosphere of *A. glacialis* and examine the influence of these AHLs on the ability of beneficial and opportunistic bacteria to colonize the phycosphere of *A. glacialis.*

## Results and discussion

### Bacterial attachment and influence on diatom physiology

*A*. *glacialis* strain A3 (deposited as CCMP3542) was isolated from the Persian Gulf and identified as previously described (Behringer et al., 2018). Axenic *A. glacialis* strain A3 cultures were generated using antibiotics as described previously (Amin et al., 2015). Under optimal growth conditions, we noticed that axenic *A. glacialis* strain A3 mostly existed as single cells or chains with an average of approximately three cells per chain while xenic *A. glacialis* strain A3 at the same cell density (~1.5×10^5^ cells/mL) formed longer chains and/or aggregated cells (Supplementary Fig. S1A and S1B). Quantifying the abundance of diatom chains spanning 1-3 cells and >3 cells per chain in axenic and xenic cultures showed that chain length did not change appreciably throughout the growth of axenic cultures with roughly half the population forming >3 cells per chain. In contrast, bacteria significantly increased the abundance of longer diatom chains, with 79.4% of the population forming >3 cells per chain relative to axenic cultures in late-exponential to early-stationary phases (Supplementary Fig. S1C). In diatoms, current evidence suggests that chain length is influenced by increasing CO_2_ concentrations (Ramos et al., 2014) and grazing pressure (Amato et al., 2018). The finding that bacteria can influence diatom chain length is novel and consistent with observations that bacteria can influence diatom cell size and morphology (Windler et al., 2014).

Removal of free-living bacteria in xenic *A. glacialis* cultures using gravity filtration through a 3-μm membrane filter and staining filtered diatom and bacterial cells with SYBR Green I showed large aggregates of bacteria on and/or in close proximity to diatom cells (Supplementary Fig. S1D). Further staining of the sample with alcian blue, a dye that stains diatom transparent exopolymeric particles (TEP), showed that bacteria mostly attach to TEP (Supplementary Fig. S1E and S1F), an observation consistent with previous observations (Bar-Zeev et al., 2012). Generating biofilm on algal TEP and/or surfaces is a typical behavior of bacteria in aquatic habitats (Kogure et al., 1981; Bagatini et al., 2014) that can enable them to persist in such environments (Geng and Belas, 2010). Forming biofilm on algal surfaces or algal TEP also protects bacteria against toxins and antibiotics and provides shelter from predation (Carvalho, 2018). For example, bacteria residing in a biofilm can tolerate antimicrobial agents at concentrations 100-1000 times needed to kill planktonic cells (Lewis, 2001). In addition, colonizing the phycosphere by attaching to TEP may help bacteria to conserve energy that would otherwise be spent on motility and chemotaxis to remain in proximity of the phycosphere (Seymour et al., 2017).

### Bacterial isolation

To examine bacterial attachment and influence on diatom chain length in the phycosphere, three bacterial strains were isolated from xenic *A. glacialis* and characterized based on 16S rRNA sequence identity as *Sulfitobacter pseudonitzschiae* F5 (Rhodobacteraceae; >99% similarity to *S. pseudonitzschiae*), *Phaeobacter* sp. F10 (Rhodobacteraceae; 97% similarity to *Phaeobacter gallaeciensis*) and *Alteromonas macleodii* F12 (Alteromonadaceae; 99% similarity to *A. macleodii*) (Supplementary Fig. S2). The *Roseobacter* group (Rhodobacteraceae) is one of the most important groups of marine bacteria that primarily colonize both biotic (e.g., phytoplankton) and abiotic surfaces (e.g., marine snow) (Gram et al., 2002; Dang et al., 2008), and can comprise up to 25% of the marine bacterial community in some regions (Wagner-Döbler and Biebl, 2006). They have been shown to form a substantial component of the *A. glacialis* microbial consortium based both on 16S rRNA amplicon sequencing (Behringer et al., 2018) and shotgun metagenomics (Shibl et al., 2020). Mining the 16S rRNA sequences of the microbial community of *A. glacialis* recovered after 20 days of isolation from the field (Behringer et al., 2018), we recovered reads that display 100% sequence identity to the 16S rRNA gene of *Phaeobacter* sp. F10 and *A. macleodii* F12 and 99% sequence identity to *S. pseudonitzschiae* F5, indicating our bacterial isolates belong to the natural population of xenic *A. glacialis* and that members of this population persist through time under laboratory culturing conditions.

### Co-culture of bacterial isolates with the diatom

To test whether these bacteria attach to *A. glacialis*, co-cultures of each bacterium with the diatom were grown in batch cultures, and growth and attachment of both partners were monitored using microscopy. When co-cultured with *A. macleodii* F12, the specific growth rate (*μ*) of the diatom did not exhibit significant changes relative to axenic controls (*μ*_axenic_ = 1.02 ± 0.05 d^−1^; *μ*_co-culture_ = 1.03 ± 0.02 d^−1^) (Supplementary Fig. S3A). Likewise, the growth of the diatom did not vary significantly when co-cultured with *Phaeobacter* sp. F10 (*μ*_axenic_ = 0.81 ± 0.02 d^−1^; *μ*_co-culture_ = 0.84 ± 0.04 d^−1^) (Supplementary Fig. S3B). In contrast, *A. glacialis* co-cultured with *S. pseudonitzschiae* F5 exhibited a 27.6% increase in *μ* relative to axenic controls (*μ*_axenic_ = 0.76 ± 0.03 d^−1^; *μ*_co-culture_ = 0.97 ± 0.03 d^−1^) (Supplementary Fig. S3C). In all co-cultures, bacteria exhibited ~3 orders of magnitude increase in cell density, indicating uptake of diatom-derived organic matter (Supplementary Fig. S3D). Surprisingly, co-cultures of *S. pseudonitzschiae* F5 with *A. glacialis* showed a significant increase in longer diatom chains than axenic cultures, similar to observations in xenic cultures (Supplementary Fig. S1C). This observation may be a byproduct of enhanced growth of *A. glacialis* with *S. pseudonitzschiae* F5 or a result of a more complex mechanism of interaction.

*Sulfitobacter pseudonitzschiae* was first isolated from cultures of the toxigenic marine diatom *Pseudo-nitzschia multiseries* obtained from the North Atlantic Ocean and the Pacific Northwest, with a model strain first coined as *Sulfitobacter* sp. SA11 (Amin et al., 2015). Subsequently, several additional *S. pseudonitzschiae* strains were isolated from the diatoms *Pseudo-nitzschia multiseries*, *Skeletonema marinoi* and *A. glacialis* originating from the Atlantic Ocean, the Swedish coast, and the Persian Gulf, respectively (Hong et al., 2015; Töpel et al., 2019). These repetitive recoveries of nearly identical bacteria (>99% 16S rRNA sequence similarity) from three genera of diatoms that originated from starkly different locations with large variations in temperature, salinity and nutrients indicate that *S. pseudonitzschiae* is a globally distributed bacterium that may be a true symbiont of diatoms. *S. pseudonitzschiae* SA11 enhances the growth rate of the diatom *P. multiseries* by 19-35% compared to axenic controls partially due to the activity of the hormone IAA, which *S. pseudonitzschiae* biosynthesizes from diatom-derived tryptophan. Both organisms also exchange organosulfur compounds and ammonia to complement each other’s metabolism (Amin et al., 2015). In this study, *S. pseudonitzschiae* F5 enhanced the growth rate of the diatom *A. glacialis* by 27.6% (Supplementary Fig. S3C). Further genome sequencing and annotation showed that *S. pseudonitzschiae* F5 also possesses three complete pathways for IAA biosynthesis from tryptophan (Supplementary Table S1), suggesting it may use the same strategy as *S. pseudonitzschiae* SA11 to enhance diatom growth.

Compared with *Sulfitobacter*, *Phaeobacter* species are known to colonize marine macro- and microalgal surfaces (e.g., *Ulva australis*, *Thalassiosira rotula*) (Rao et al., 2006; Thole et al., 2012). *P. inhibens* has been shown to control bacterial community assembly in the phycosphere of *T. rotula* (Majzoub et al., 2019). *P. inhibens* also produces the hormone IAA to promote the growth of the coccolithophore, *Emiliania huxleyi*, similar to *S. pseudonitzschiae* (Segev et al., 2016). *P. gallaeciensis* has been shown to lyse senescent *E. huxleyi* by producing algicidal compounds known as roseobacticides (Seyedsayamdost et al., 2011). Although *Phaeobacter* sp. F10 did not enhance the growth rate of *A. glacialis* (Supplementary Figure S3B), metatranscriptomic analysis of a near-identical metagenomically assembled genome from the *A. glacialis* strain A3 microbial consortium indicates that *Phaeobacter* sp. F10 is also a symbiont of *A. glacialis* (Shibl et al., 2020).

Alteromonadaceae are widespread marine opportunistic copiotrophs (López-Pérez et al., 2012) that display algicidal activities with phytoplankton during algal blooms (Mayali and Azam, 2004). The *Pseudoalteromonas* and *Alteromonas* genera are known to effectively metabolize the organic matrix surrounding diatom frustules, exposing the silica shell to increased dissolution by the surrounding water (Bidle and Azam, 2001), and show strong algicidal activity by releasing dissolved substances (Mayali and Azam, 2004). For example, *A. colwelliana* shows growth inhibition and algicidal activity against the diatom *Chaetoceros calcitrans* (Kim et al., 1999). *A. macleodii* have been shown to degrade a variety of algal TEP and exopolysaccharides (Koch et al., 2019) that may enable them to benefit from phytoplankton-derived carbon without contributing to phytoplankton metabolism. In addition, *A. macleodii* has been shown to compete for nitrate with the diatom *Phaeodacylum tricornutum* in the presence of organic carbon (Diner et al., 2016).

### Bacterial attachment in the phycosphere

In order to test which bacterial strains attach to the diatom or TEP, we removed free-living bacterial cells from co-cultures of each strain with the diatom using gravity filtration through a 3-μm membrane filter and stained TEP with alcian blue, and diatom and bacterial nucleic acids with SYBR Green I (Fig. 1). Both symbiotic strains, *S. pseudonitzschiae* F5 and *Phaeobacter* sp. F10, displayed strong attenuation onto filters, while no attached *A. macleodii* F12 were observed (Fig. 1D-1F). The composite images of bright field and fluorescence showed a strong attachment preference to diatom TEP of *S. pseudonitzschiae* F5 and *Phaeobacter* sp. F10 in co-cultures with *A. glacialis* as observed in xenic cultures (Fig. 1G-1H). Surprisingly, *A. macleodii* F12 did not show any attachment capacity to *A. glacialis* or TEP (Fig. 1) despite its ability to degrade algal polysaccharides (Koch et al., 2019). These observations suggest an inherent mechanism that enables the roseobacters but not *A. macleodii* to attach to TEP. To shed more light on such mechanisms, we sequenced the genomes of all three isolates.

**Fig. 1.**
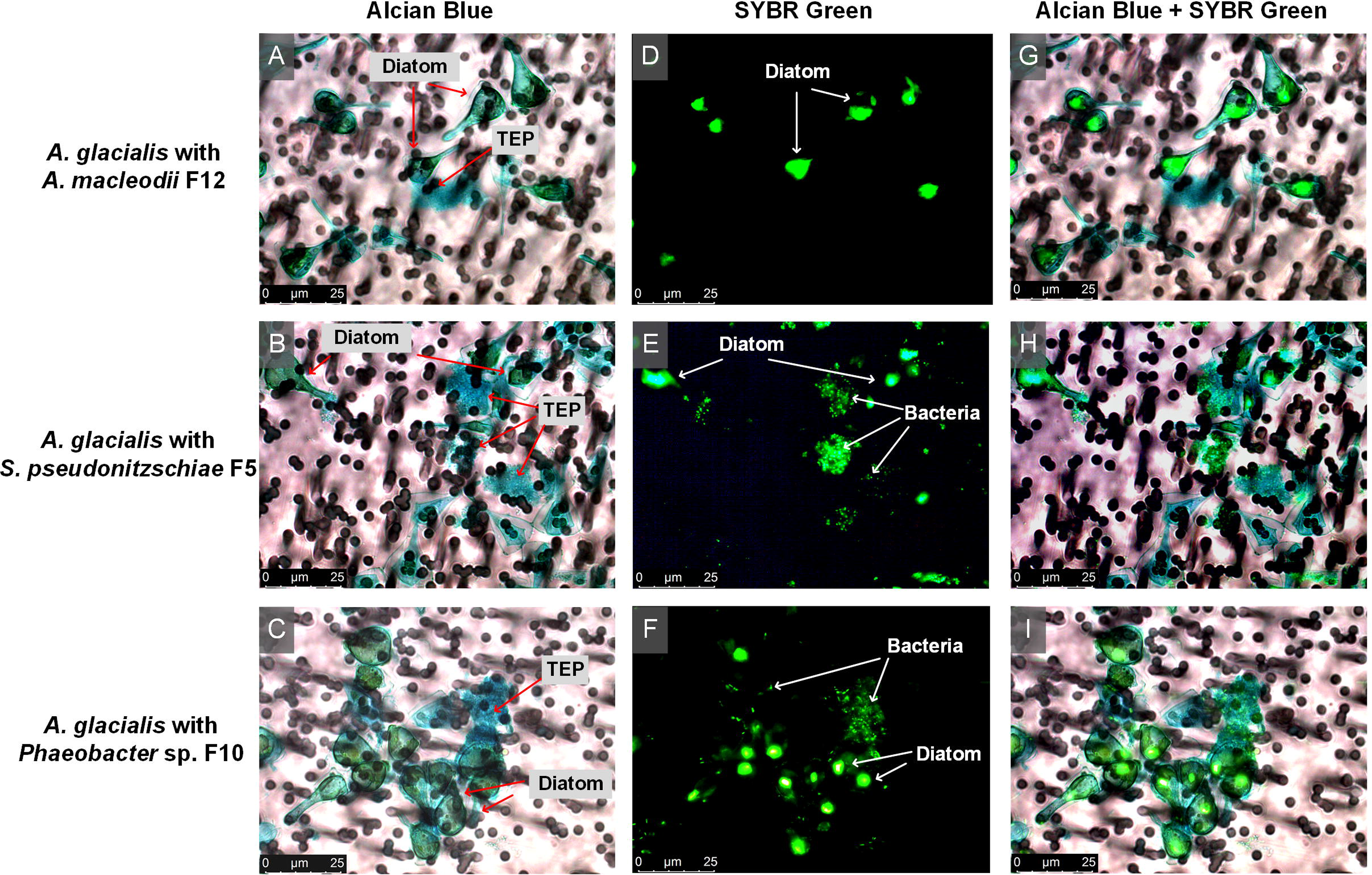
Micrographs of *A. glacialis* strain A3 co-cultures with bacterial isolates. Diatom TEP were stained with alcian blue **(A, B, C)**. Diatom and bacterial DNA was stained with SYBR Green I **(D, E, F)**. Composite images show bacterial attachment mostly to TEP for *S. pseudonitzschiae* F5 and *Phaeobacter* sp. F10 but not *A. macleodii* F12 **(G, H, I)**. Background in light micrographs show membrane filters.

### Genomic comparisons and AHLs identification

The genomes of the three strains were obtained by a combination of PacBio and Illumina sequencing as described in the methods. The GC contents of both *Roseobacter* group member genomes were more similar to each other (61.8% for *S. pseudonitzschiae* F5 and 60.0% for *Phaeobacter* sp. F10), while *A. macleodii* F12 had a lower GC content (44.9%) (Table 1), consistent with the phylogenetic similarities of *S. pseudonitzschiae* F5 and *Phaeobacter* sp. F10 compared to *A. macleodii* F12 (Supplementary Fig. S2). *S. pseudonitzschiae* F5 possessed the largest estimated genome size (5.1 Mb) compared to *Phaeobacter* sp. F10 (4.0 Mb) and *A. macleodii* F12 (4.7 Mb) (Table 1). Consistent with this observation, *S. pseudonitzschiae* F5 also had the greatest number of predicted genes (4991) compared with either *Phaeobacter* sp. F10 (3878) or *A. macleodii* F12 (4654) (Supplementary Fig. S4A). A major difference between the three genomes was the apparent higher number of putative genes involved in membrane transport of substrates, particularly primary active transporters (e.g., amino acids, sugars) (Mishra et al., 2014). For example, the relative abundance of putative membrane transporters in the *S. pseudonitzschiae* F5 genome is 10.50% compared to *Phaeobacter* sp. F10 (8.44%) or *A. macleodii* F12 (7.05%) when normalized to genome size, with primary active transporters being the most abundant in the roseobacters’ genomes but not in *A. macleodii* F12 (Supplementary Fig. S4B). This observation suggests that both symbionts are more attuned to phycosphere metabolites than *A. macleodii* F12.

**Table 1.** Bacterial genome statistics.

| Statistic | *S. pseudonitzschiae* F5 | *Phaeobacter* sp. F10 | *Alteromonas macleodii* F12 |
| --- | --- | --- | --- |
| Genome size (bp) | 5,087,289 | 4,008,202 | 4,727,977 |
| Number of contigs | 17 | 3 | 5 |
| Chromosome size (bp) | 3,764,450* | 3,783,964 | 3,117,849 |
| CDs | 4,991 | 3,878 | 4,654 |
| GC content (%) | 61.8 | 60.0 | 44.9 |
| N50 value | 179,155 | 153,588 | 3,122,272 |
| L50 value | 3 | 1 | 1 |

\**S. pseudonitzschiae* F5 chromosome size estimation was not possible due to high level of repeats in the genome. The size indicated is the size of the largest contig.

Chemotaxis, motility, and attachment presumably contribute to successful colonization of phytoplankton surfaces. To examine the ability of all three bacteria to interact with the diatom phycosphere, we conducted a genome-wide analysis to compare genes involved in bacterial chemotaxis, motility, attachment, and quorum sensing. All three genomes contained four chemotaxis genes in a single operon-like structure (*cheA*, *cheR*, *cheW*, and *cheY*), while a fifth gene, *cheB*, was present in *Phaeobacter* sp. F10 and *A. macleodii* F12, but absent in *S. pseudonitzschiae* F5 (Fig. 2A and Supplementary Table S2). *CheAYW* together mediate a signal transduction cascade that functions to regulate flagellar motors, while *cheB* and *cheR* together regulate the methylation state of methyl-accepting chemotaxis proteins (Wuichet et al., 2007). Flagellar genes were present in all three genomes, although *S. pseudonitzschiae* F5 and *Phaeobacter* sp. F10 also contained a suite of flagellar structure genes absent in *A. macleodii* F12 (*flgN*, *fliG*, *flgJ*, *motB*, and MotA/TolQ/ExbB proton channel family protein). In addition, both *S. pseudonitzschiae* F5 and *Phaeobacter* sp. F10 contained 11 genes related to pilus formation in contrast to *A. macleodii* F12, which only contained a pilin *flp* gene, which is involved in pilus formation (Bardy et al., 2003). For attachment, genes involved in exopolysaccharide production and biofilm formation were distributed throughout the genomes of the three bacteria (Fig. 2A and Supplementary Table S2). These lines of evidence suggest that *S. pseudonitzschiae* F5, *Phaeobacter* sp. F10, and *A. macleodii* F12 have the ability to carry out chemotaxis, construct flagella and pili structures, and form biofilm. Despite this observation, *A. macleodii* F12 did not attach to *A. glacialis* or TEP produced by *A. glacialis* (Fig. 1).

**Fig. 2.**
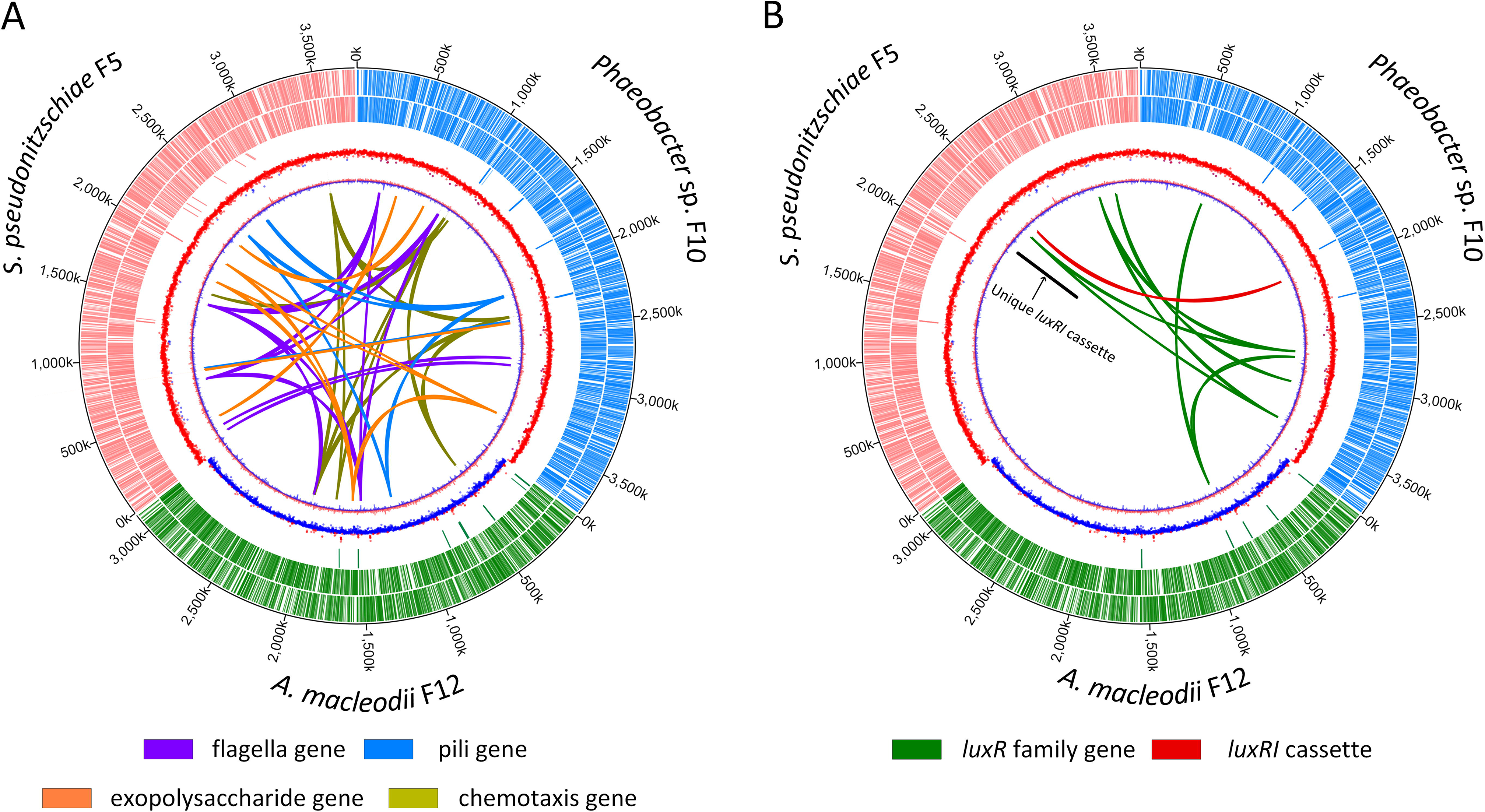
Chromosomal maps of bacterial isolate genomes showing distribution and localization of loci involved in attachment, motility, chemotaxis and quorum sensing. Tracks from the outermost to the center represent position on each chromosome, forward-strand protein-coding genes, reverse-strand protein coding genes, rRNA and tRNA genes, GC content and GC skew, respectively. Chromosomes of *S. pseudonitzschiae* F5, *Phaeobacter* sp. F10, and *A. macleodii* F12 are color-labeled with red, blue and green, respectively. **(A)** Homologs related to chemotaxis, motility and attachment. **(B)** Homologs related to quorum sensing. Black line refers to a *luxI* gene homolog unique to *S. pseudonitzschiae* F5.

Quorum sensing (QS) autoinducers (e.g., AHLs) have been widely shown to modulate important biological functions, such as biofilm formation and motility (Bassler, 2002; Waters and Bassler, 2005), indicating QS may be able to regulate bacterial ‘swim-or-stick’ lifestyles in the phycosphere and suggesting that *A. macleodii* F12 may lack a functioning QS system. Indeed, *luxI*-like genes that are responsible for biosynthesizing AHLs were only present in *S. pseudonitzschiae* F5 (2 homologs) and *Phaeobacter* sp. F10 (1 homolog), while *A. macleodii* F12 completely lacked apparent *luxI* homologs (Fig. 2B and Supplementary Table S2). Indeed, *luxI* is rarely present in *Alteromonas* species. We found only two of 67 genomes belonging to the Alteromonadaceae that are publicly available contain putative AHL synthases. In addition to AHL synthesis, a transcriptional regulator, encoded by a *luxR* gene, is required to perceive AHLs and coordinate gene expression among bacterial populations. All three genomes contained *luxR* family genes, while *S. pseudonitzschiae* F5 possessed four putative *luxR*-family genes, *Phaeobacter* sp. F10 possessed six and *A. macleodii* F12 possessed only one (Fig. 2B and Supplementary Table S2). Typically, apparent *luxR* genes are found adjacent to or near a *luxI* gene on bacterial chromosomes in a single operon-like structure, such as in *S. pseudonitzschiae* F5 and *Phaeobacter* sp. F10 (Supplementary Table S2). However, some bacteria also have putative ‘solo’ *luxR* genes without associated putative *luxI* genes, which is the case in *A. macleodii* F12 and for the additional putative *luxR* homologs present in *S. pseudnonitzschiae* F5 and *Phaeobacter* sp. F10. It has been hypothesized that bacteria possessing ‘solo’ *luxR* genes do so to detect and respond to exogenous signals from other bacterial populations (Hudaiberdiev et al., 2015) or that these solo genes represent the loss of QS function (Subramoni and Venturi, 2009). Some solo LuxR proteins have low specificity for AHL binding compared with LuxR proteins coupled with LuxIs (Subramoni and Venturi, 2009). For example, SdiA, a solo transcriptional regulator that is present in members of *Salmonella*, *Escherichia*, and *Klebsiella*, could bind seven different AHL molecules (Yao et al., 2006; Janssens et al., 2007). Potentially, the solo *luxR* genes in *A. macleodii* F12 and the symbionts may respond to a wide range of AHLs from other bacteria.

To characterize AHLs produced by *S. pseudonitzschiae* F5 and *Phaeobacter* sp. F10, we purchased a suite of commercially available AHL standards (Supplementary Table S2) and used ultra-high-performance liquid chromatography-tandem mass spectrometry (UHPLC-MS/MS) to identify potential AHLs in bacterial cultures. Three AHLs were recovered from *S. pseudonitzschiae* F5 and *Phaeobacter* sp. F10 pure culture supernatants with ionized parent masses of m/z 270.1700, 352.2480, and 354.2640 and identified as 3-oxo-C_10_-HSL, 3-oxo-C_16:1_-HSL and 3-oxo-C_16_-HSL, respectively, using high-resolution m/z values, daughter-ion fragmentation and retention times (Fig. 3 and Supplementary Fig. S5). No standard from the AHLs library matched any metabolite in the supernatant of *A. macleodii* F12, consistent with its lack of *luxI*-like genes (Fig. 2). We attempted to measure AHLs in co-cultures of the roseobacters and the diatom but due to the lower bacterial abundance in cocultures compared to pure bacterial cultures we were not successful in detecting AHLs. The identification of AHL production by diatom symbionts prompted examination of the influence of these signaling molecules on the ability of the symbionts to attach to diatom TEP.

**Fig. 3.**
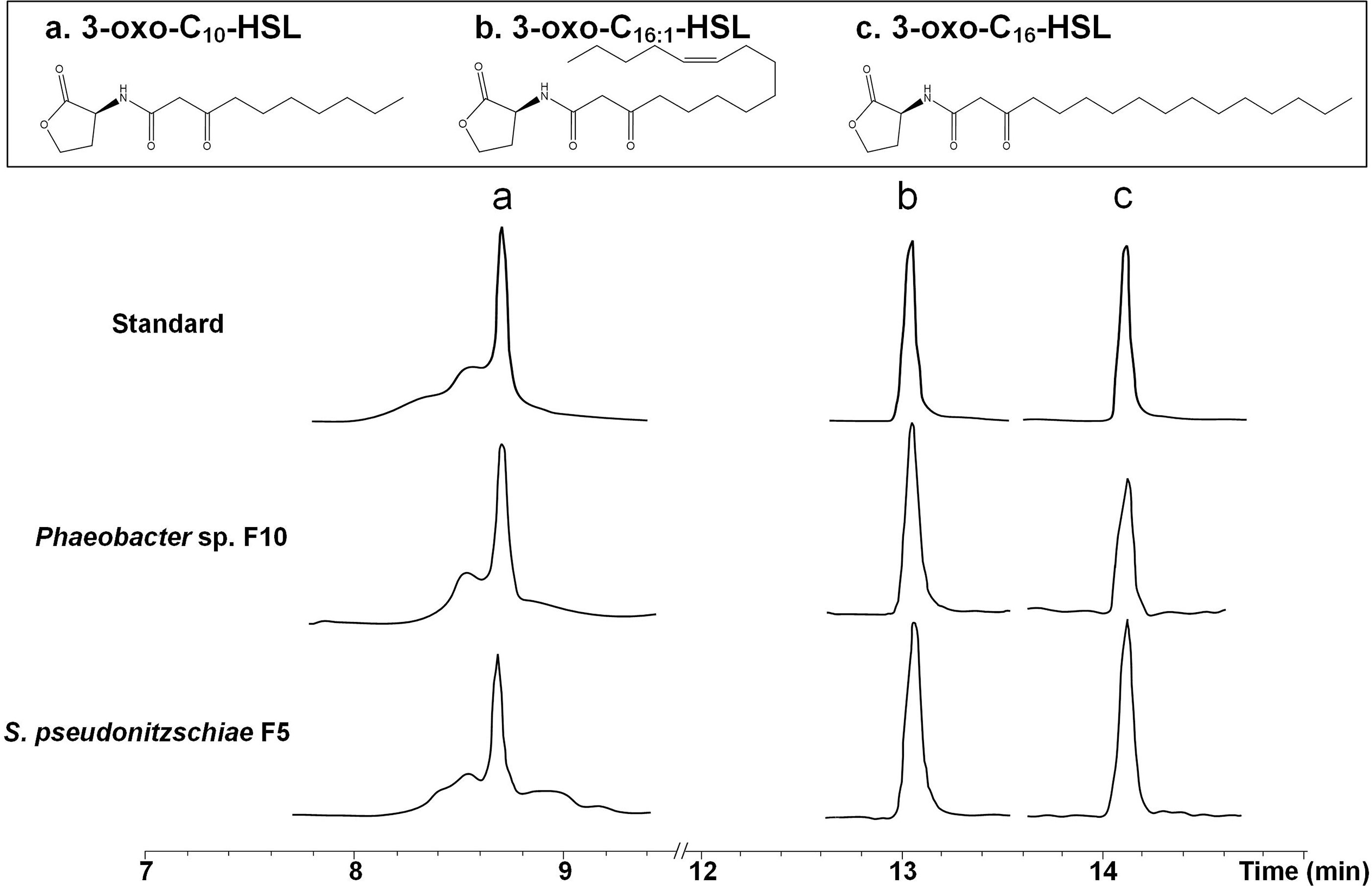
Identification of acyl-homoserine lactones (AHLs) from roseobacters isolates. Structures and UHPLC–MS/MS chromatograms of AHLs of *S. pseudonitzschiae* F5 or *Phaeobacter* sp. F10 isolated from pure cultural supernatants compared to purchased standards. For complete MS/MS spectra refer to Supplementary Fig. S5.

### Influence of AHLs on bacterial motility and biofilm formation

Cell attachment is the first step in the process of biofilm formation, which requires motility for initial attachment to surfaces (Slightom and Buchan, 2009). Consequently, we examined the influence of AHLs on motility and biofilm formation since both functions are regulated by QS in bacteria (Hammer and Bassler, 2003; Daniels et al., 2004). All three AHLs and two QS inhibitors (QSIs), 2(*5H*)-furanone and furanone C-30 (Ponnusamy et al., 2010; He et al., 2012), were used to test the motility and biofilm formation capabilities of *S. pseudonitzschiae* F5, *Phaeobacter* sp. F10, and *A. macleodii* F12. QSIs were used to confirm that any phenotypes observed using AHLs are due to QS regulation and not byproducts of other processes. Since the *A. macleodii* F12 genome contained a putative solo *luxR* gene (Fig. 2B), we suspected *A. macleodii* F12 might still respond to AHLs produced by *S. pseudonitzschiae* F5 and *Phaeobacter* sp. F10 despite lacking the ability to synthesize AHLs.

The addition of 2 μM of each AHL to each bacterium showed statistically significant inhibition of motility in *S. pseudonitzschiae* F5 and *Phaeobacter* sp. F10 (p<0.01) by all three AHLs, while both QSIs enhanced motility in *S. pseudonitzschiae* F5, as expected, and furanone C-30 enhanced motility in *Phaeobacter* sp. F10 (Fig. 4A and 4B). Specifically, 3-oxo-C_16:1_-HSL exhibited the strongest inhibition of motility in *S. pseudonitzschiae* F5 by 78.2% (p<0.001) compared to the two other AHLs (28.6% and 31.8%). Despite showing two AHLs, 3-oxo-C_16_-HSL and 3-oxo-C_10_-HSL, weakly inhibiting motility in *A. macleodii* F12, both QSIs also weakly inhibited its motility, suggesting *A. macleodii* F12 does not respond or weakly responds to QS molecules (Fig. 4A and 4B). Bacterial motility is characterized by a circular swimming zone in the motility assay or by dendritic morphology, which is typical for swarming motility caused by a locally restricted movement on agar plates (Michael et al., 2016). *Phaeobacter* sp. F10 and *A. macleodii* F12 displayed a typical circular swimming zone while *S. pseudonitzschiae* F5 displayed a dendritic motility phenotype (Fig. 4A).

**Fig. 4.**
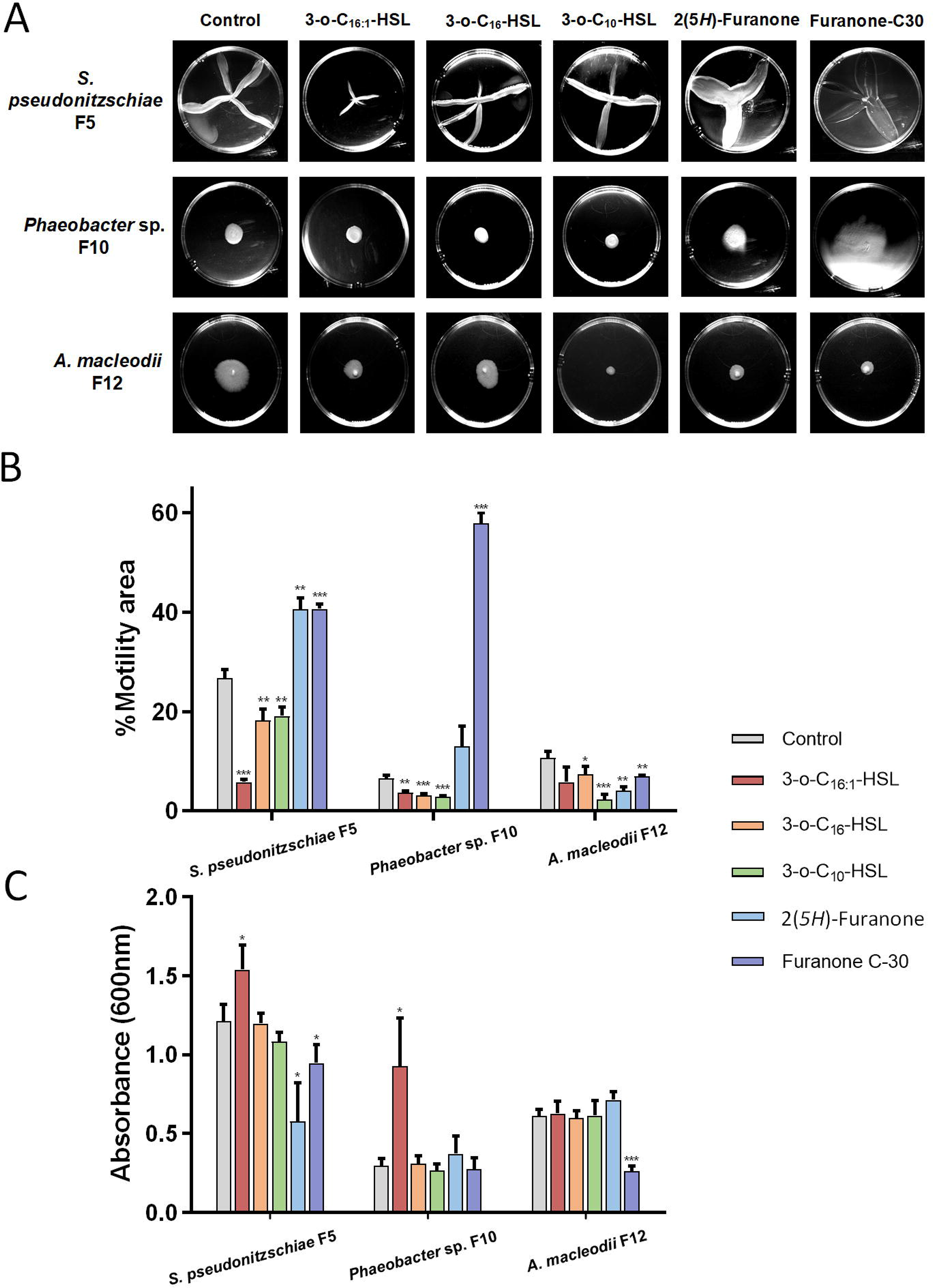
AHLs control bacterial motility and attachment in *Roseobacter* group bacteria. **(A)** Motility assays of *S. pseudonitzschiae* F5, *Phaeobacter* sp. F10 and *A. macleodii* F12 with three AHL molecules: 3-oxo-C_10_-HSL, 3-oxo-C_16_-HSL and 3-oxo-C_16:1_-HSL, and two quorum sensing inhibitors (QSIs): 2(*5H*)-furanone and furanone C-30. Assays were conducted by inoculating 0.25% soft agar plates with each bacterium and incubating for three days. **(B)** Percent Motility of *S. pseudonitzschiae* F5, *Phaeobacter* sp. F10 and *A. macleodii* F12 in the presence of 2 μM AHLs or QSIs. **(C)** Biofilm quantification in the presence of 10 μg/mL AHLs or QSIs. Assays were conducted by incubating 1×10^6^ bacterial cells with each molecule in 96-well plates for 24 hours. Attachment was quantified using absorbance of solubilized crystal violet at 600 nm. All error bars represent standard deviation (S.D.) of triplicate experiments. * p<0.05, ** p<0.01, *** p<0.001).

To assess biofilm formation, a standardized crystal violet assay was conducted as described previously (O’toole and Kolter, 1998). Interestingly, 10 μg/mL of 3-oxo-C_16:1_-HSL significantly enhanced biofilm formation in *S. pseudonitzschiae* F5 and *Phaeobacter* sp. F10 by 26.7% and 211%, respectively (p<0.05), while 3-oxo-C_10_-HSL and 3-oxo-C_16_-HSL did not influence biofilm formation in either bacteria (Fig. 4C). These results suggest each AHL molecule regulates different sets of functions. Similar observations have been shown previously (Su et al., 2018; Hou et al., 2019). In contrast, both QSIs inhibited biofilm formation of *S. pseudonitzschiae* F5 by 52.2% and 21.9%, respectively, relative to the control. None of the AHLs influenced biofilm formation of *A. macleodii* F12 (Fig. 4C).

These findings suggest that specific AHLs produced by diatom symbionts promote bacterial colonization of the phycosphere by enhancing bacterial capacity to form biofilms and reduce motility, both are functions essential to successfully colonize the phycosphere. In contrast, *A. macleodii* F12 is unable to respond to these molecules or to effectively attach to TEP. Other mechanisms must also be functioning to further prevent opportunists, like *A. macleodii* F12, from benefitting from diatom exudates. One such mechanism is the regulation of microbial consortia by the eukaryotic host, *A. glacialis*. *A. glacialis* has recently been shown to release the unusual metabolites, rosmarinic acid and azelaic acid, in the phycosphere in response to bacteria (Shibl et al., 2020). Azelaic acid promoted the growth of both symbionts and concomitantly inhibited the growth of *A. macleodii* F12. Similarly, rosmarinic acid inhibited the motility of *S. pseudonitzschiae* F5 and *Phaeobacter* sp. F10 in an identical way to 3-oxo-C_16:1_-HSL, while upregulating motility in *A. macleodii* F12 (Shibl et al., 2020). Rosmarinic acid has been recently shown to interfere with QS in a plant pathogen (Hammer and Bassler, 2003), suggesting this molecule acts as a QSI. A similar mechanism of interfering with QS, a process known as quorum quenching, is used by the marine macroalga *Delisea pulchra* to inhibit swarming motility of *Serratia liquefaciens* by producing halogenated furanones (Rasmussen et al., 2000). Cumulatively, our observations suggest that *S. pseudonitzschiae* F5 and *Phaeobacter* sp. F10 switch their free-living mode to surface-attached mode to remain in the phycosphere by releasing AHLs. Combined with the diatom host release of unique secondary metabolites that further promote symbiont phycosphere colonization and inhibit colonization of opportunists, these mechanisms ensure diatoms are not prey to random encounters with opportunists and pathogens.

### QS gene homology and organization in the *Roseobacter* group

To shed light on the prevalence of QS systems in the *Roseobacter* group, we constructed a phylogenetic tree of 52 sets of the LuxR-ITS-LuxI sequences (*luxRI* cassettes) in 32 *Roseobacter* group representative genomes. The tree revealed multiple conserved QS gene topologies that were distributed across the *Roseobacter* group (Fig. 5).

**Fig. 5.**
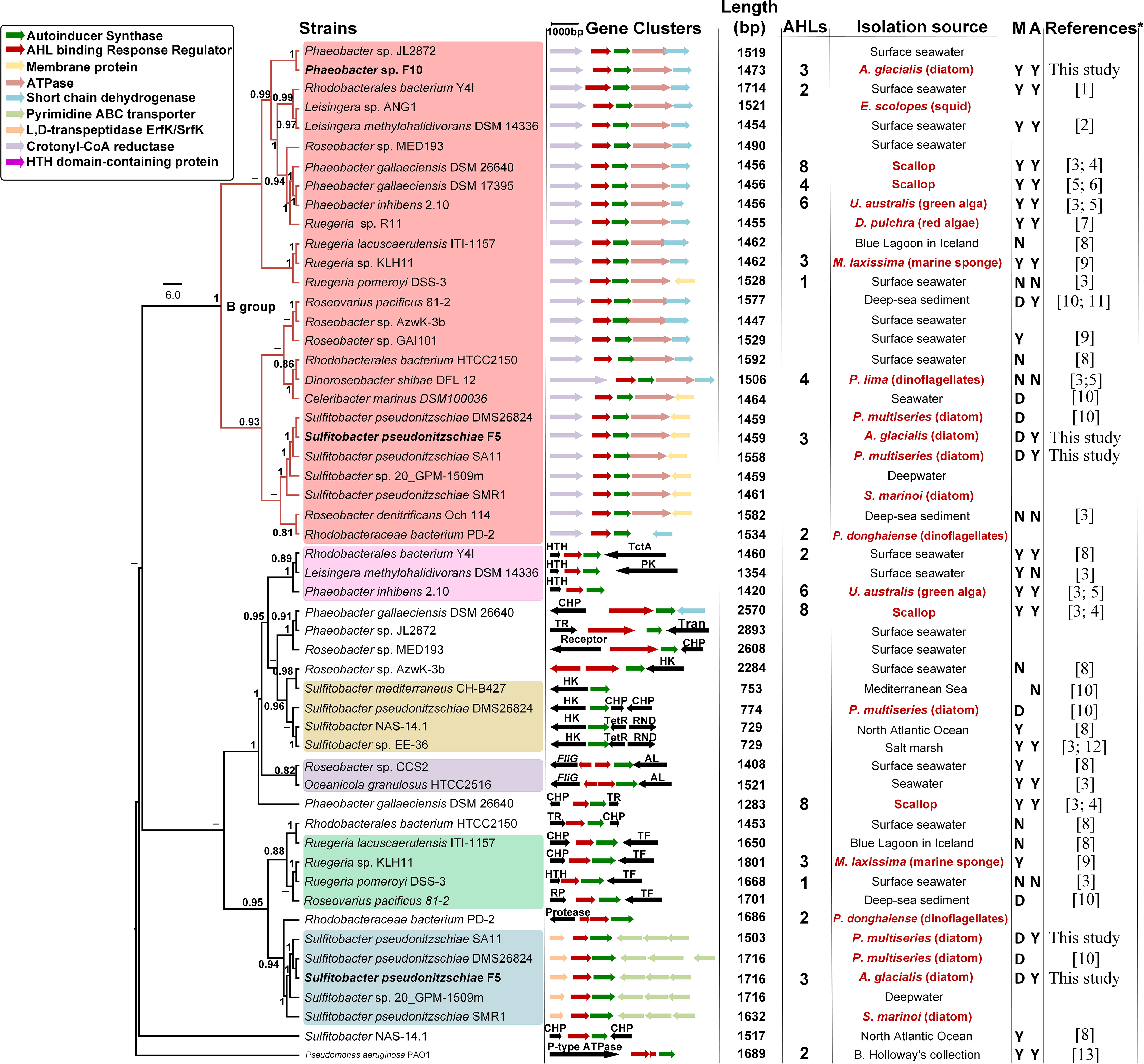
Maximum-likelihood phylogenetic tree of 50 LuxR-ITS-LuxI loci and neighboring genes in 32 roseobacterial genomes. Group B is highlighted by red branches at the top of the tree. Color shades indicate the different groups of luxR/I cassettes. Bacteria isolated in this study are bold-faced. Length shows the sequence length of LuxR-ITS-LuxI loci. D = dendritic motility, M = motility, A = attachment. Y and N indicate whether a bacterium is able or unable to carry out each function. Isolation origin of bacteria highlighted in red indicate a biological host origin. The number of AHL molecules reported from each bacterium is provided, if known. Note that information on which cassette makes which AHL molecule is scarce. Some bacteria are present twice in the tree because they possess more than one Lux-like operon. Gene neighborhood abbreviations: HK, histidine kinase; CHP, conserved hypothetical protein; TctA, TctA family transmembrane transporter; RND, RND multidrug efflux pump; AL, adenylosuccinate lyase; TF, trigger factor; Tran, transposase; TetR, TetR family transcriptional regulator; TR, transcriptional regulator. Bootstrap values >50% calculated from 1000 iterations are shown at branch nodes. *Full references are listed in Supplementary Information.

In Figure 5, half of the roseobacters genomes contained more than one pair of *luxRI* (18/31) and more than two AHL molecules were identified in nine strains. It has been shown that >80% of bacteria in the *Roseobacter* group possess at least one *luxRI* gene cassette (Buchan et al., 2016). *Phaeobacter* species consistently appear to produce more than three AHLs using 2−3 *luxI* homologs (Fig. 5 and Supplementary Table S3). For example, *Phaeobacter gallaeciensis* DSM 26640 produces up to 8 different AHLs via only three *luxI* homologs (Ziesche et al., 2015). Cude and Buchan (2013) provide a classification of the genetic makeup and organization of the *luxRI* gene cassettes and neighboring genes, with the B topology (B group) being the most common cassette structure (Fig. 5). In our work, B group contained 26 *luxRI* cassettes distributed in 26 genomes (Fig. 5). Both *S. pseudonitzschiae* F5 and *Phaeobacter* sp. F10 possess one cassette that belongs to this group, while a unique *luxRI* cassette only occurs in *S. pseudonitzschiae* F5. Interestingly, both *S. pseudonitzschiae* F5 and *Phaeobacter* sp. F10 strains produced the same three AHLs molecules (Fig. 3 and Supplementary Fig. S5), suggesting these molecules must be a byproduct of a homologous *luxI*, common to both *S. pseudonitzschiae* F5 and *Phaeobacter* sp. F10. This finding is consistent with previous studies showing that a single *luxI*-type gene is responsible for biosynthesizing more than one AHL molecule (Ortori et al., 2007; Hansen et al., 2015). LuxI proteins use S-adenosylmethionine (SAM) and acylated acyl-carrier protein (ACP) to catalyze the acylation and lactonization of AHL molecules (Churchill and Chen, 2011). Potentially, the *luxI* homolog common to *S. pseudonitzschiae* F5 and *Phaeobacter* sp. F10 does not discriminate well between different substrates and can accept acyl chains of varying lengths, enabling it to biosynthesize three AHLs. This lack of specificity could allow different AHL molecules produced by a single protein to bind different transcriptional regulators and thus regulate different functions. By regulating the availability of substrates needed to make each AHL, different AHLs may be produced depending on the environmental condition while continuously expressing the same AHL synthase. For example, different temperatures have been shown to influence the types and concentrations of seven different AHLs produced by one *luxI* in the fish pathogen *Aliivibrio salmonicida* (Hansen et al., 2015). The diversity of substrates and selectivity levels of different *luxI* homologs also influence the proportions of various AHLs produced by roseobacters (Ziesche et al., 2018). *S. pseudonitzschiae* F5 and *Phaeobacter* sp. F10 were isolated from the same diatom, indicating horizontal gene transfer may be responsible for their identical production of AHLs. The second *luxI* gene in *S. pseudonitzschiae* F5 likely produces one or more other AHL molecules that we were not able to characterize due to the limitation of AHL mass spectrometry standards. As mentioned before, *S. pseudonitzschiae* F5 displayed a dendritic motility phenotype, which is absent in most roseobacters (Bartling et al., 2018). It is not clear what ecological advantage dendritic motility have on bacteria like *S. pseudonitzschiae*. Interestingly, two additional *S. pseudonitzschiae* strains displayed this phenotype previously (Bartling et al., 2018), indicating this phenotype is common in this species for unknown reasons.

The phycosphere is a unique environment that enables the accumulation of organic molecules in close proximity to phytoplankton cells along with their associated bacterial populations. Mounting evidence indicates that DOM secretions by phytoplankton lead to the accumulation of higher bacterial density in the phycosphere when compared to bulk seawater (Blackburn et al., 1998; Smriga et al., 2016), which is the ideal scenario under which QS systems are activated. The isolation sources of bacteria in Figure 5 indicate that 37.5% of strains were isolated from a eukaryotic host and 21.9% from phytoplankton and that these roseobacters produce a wide variety of long-chain AHLs (C_10_-C_18_) (Supplementary Table S3). Long-chain AHLs are more stable in alkaline environments, such as the phycosphere, compared to short-chain AHLs (Yates et al., 2002). Thus, the production of long-chain AHLs by roseobacters in the phycosphere may lead to a higher local concentration than short-chain AHLs, and thus enable roseobacters to respond collectively to the phycosphere environment more successfully than other bacteria.

Here, we have shown that two diatom symbionts use AHL molecules to inhibit their motility and enhance biofilm formation, processes that likely control their ability to attach to diatom TEP and thus colonize the phycosphere. In contrast, an opportunist bacterium was incapable of attaching to diatom TEP and was found to lack the ability to synthesize AHL molecules. The advantage of symbionts to switch their lifestyle from motile bacteria entering the phycosphere to permanent residents of this microenvironment is essential to marine bacteria that rely on phycosphere exudates to survive. This ‘swim-or-stick’ switch appears also to be partially controlled by diatoms that can make quorum sensing mimics, such as rosmarinic acid. Significant work is needed to delineate the importance of bacterial intraspecies signaling vs eukaryotic host interference in bacterial communication.

## Experimental procedures

### Diatom growth

*Asterionellopsis glacialis* strain A3 was deposited for this work in the National Center for Marine Algae and Microbiota (NCMA) under the accession number CCMP3542. Axenic *A. glacialis* strain A3 (CCMP3542) was generated as described previously (Behringer et al., 2018). All diatom cultures were grown at 22℃ in a 12:12 hour light/dark diurnal cycle (125 μE m^−2^ s^−1^) in semi-continuous batch cultures (Brand et al., 1981). Since *in vivo* fluorescence linearly correlates with cell numbers during exponential growth in batch cultures (Wood et al., 2005), diatom growth was monitored by measuring *in vivo* fluorescence of chlorophyll *a* (relative fluorescence units, RFU) using a 10-AU fluorimeter (Turner Designs, San Jose, CA, United States). Specific growth rates (*μ*) of diatoms were calculated from the linear regression of the natural log of RFU versus time during the exponential growth phase of cultures. The standard deviation of *μ* was calculated using values from biological replicates (*n* = 5 unless otherwise indicated) over the exponential growth period. The relationship between cell density and RFU of *A. glacialis* strain A3 can be formulated with the following equation: y = 14.958x - 0.519, where y is cells/μL and x is the RFU value, calculated from the regression line of linear portion of the growth curve. The adjusted R^2^ value for this curve is 0.98 (p < 0.001).

### Bacterial isolation, identification and phylogenetic analysis

To assess whether specific bacteria attach to *A. glacialis* strain A3, we cultivated bacteria from the xenic *A. glacialis* strain A3 culture as described previously (Shibl et al., 2020). Briefly, bacteria were isolated from the diatom at early stationary phase by serially diluting 0.5 mL 1000 times in sterile f/2 media (Guillard, 1975) and subsequently spreading 200 μL onto (1) marine agar plates, (2) plates containing per liter of seawater: 15g agar and 2g carbon source (sodium succinate or glucose) and (3) sterilized *A. glacialis* strain A3 culture suspension with 1.5% agar. Plates were incubated at 25℃ in the dark for 3-7 days and single colonies with unique morphologies were re-streaked 3 times to eliminate cross-contamination before storage at −80℃ in 15% glycerol stocks.

Bacterial isolates were identified using direct PCR (Hofmann and Brian, 1991). Isolates were incubated in 3 mL marine broth at 25℃ in the dark until optical density (600 nm) reached 1.0; subsequently, 2 μL of each bacterium was used as a DNA template in PCR. The 16S rRNA gene from all bacteria was amplified by universal primers (27F, 1492R) as previously described (Amin et al., 2015). PCR products were purified by the Wizard PCR purification kit (Promega) and sequenced using Sanger sequencing (Apical Scientific, Malaysia). Sequences were aligned with 16S rRNA sequences from GenBank using ClustalW Multiple Alignment in BioEdit 7.0.9.0. A neighbor-joining (NJ) phylogenetic tree was constructed with BIONJ (Gascuel, 1997) using Kimura’s two-parameter model. Another tree was constructed using the Maximum Likelihood (ML) methods using PhyML 3.0 (Guindon et al., 2010) with GTR+R+I substitution model as determined by SMS (Lefort et al., 2017). The final 16S rRNA phylogenetic consensus tree was generated and edited using FigTree 1.4.2 (Rambaut, 2014). 16S rRNA of strains *S. pseudonitzschiae* F5, *Phaeobacter* sp. F10, and *A. macleodii* F12 were compared with the microbial community of *A. glacialis* recovered after 20 days of isolation from the field (Behringer et al., 2018) using the BLASTn tool in BLAST+ (Camacho et al., 2009).

### Co-culture generation and growth

For co-cultures, axenic *A. glacialis* strain A3 was inoculated from an acclimated, mid-exponential phase growing culture into 25 mL sterile f/2 media to achieve an initial diatom cell density of ~4,000 cells/mL. Bacteria were grown in marine broth (ZoBell, 1941) in the dark overnight from a single colony at 26℃ and shaking at 180 rpm. Cultures were centrifuged at 4000 rpm for 10 min followed by washing twice with sterile f/2 medium. Subsequently, cultures were used to inoculate axenic *A. glacialis* cultures at an initial bacterial density of ~1×10^4^ cells/mL. Bacterial counts in co-cultures were quantified by staining 1 mL fresh bacterial cells with 1x SYBR Safe stain (Edvotek Corp. USA), incubating stained samples in the dark at room temperature for one hour and using a CyFlow Space flow cytometer (Partec, Münster, Germany). Specific growth rates of diatoms were calculated as described above. Specific growth rates (*μ*) of bacteria were calculated from the linear regression of the natural log of cell counts from 2-6 days of cultures. The standard deviation of *μ* was calculated using values from biological replicates (*n* = 3 unless otherwise indicated).

### Bright-field and fluorescence microscopy

To observe transparent exopolymeric particles (TEP) distribution and their attached bacteria, two staining steps were applied. Alcian blue was first used to stain TEP in diatom cultures. 1 mL culture in mid-exponential phase was gently filtered by gravity onto 3-μm 25 mm polycarbonate membrane filters (Whatman) to remove free-living bacteria (~ 1 μm). Subsequently, 1 mL alcian blue in 0.06% glacial acetic acid (pH 2.5) was allowed to gently run down the tube wall onto the filter to stain samples for 10 min at room temperature (Long and Azam, 1996). Excess alcian blue was removed using a gentle wash with PBS buffer and gently filtered by gravity. Subsequently, filters that only contained diatoms (~25 μm for a single cell), TEP particles and attached bacteria were fixed with Moviol-Sybr Green I as described previously (Lunau et al., 2005). Filters were visualized using a DMI6000B epifluorescence microscope (Leica) with a DMC2900 color brightfield camera (Leica). Bright-field microscopy was used for TEP observation. Fluorescence microscopy was used for bacteria and diatom nucleic acid observations. L5 fluorescence filter was used for SYBR green I staining of both bacterial and algal nucleic acids with an excitation wavelength range of 480 - 656 nm and an emission wavelength at 590 nm. Y5 fluorescence filter was used for algal chlorophyll autofluorescence with the excitation wavelength of 620 nm and an emission wavelength of 700 nm. Both bright-field and fluorescence images were merged using LAS X software (Leica Microsystems, Germany). The chain length of axenic, xenic *A. glacialis* strain A3 and co-culture of the diatom with *S. pseudonitzschiae* F5 were quantified using an inverted microscope (Eclipse Ti-U, Nikon, Japan).

### Bacterial genome sequencing and assembly

Genomic DNA of *S. pseudonitzschiae* F5, *Phaeobacter* sp. F10, and *A. macleodii* F12 were extracted from pure bacterial cultures using an E.Z.N.A. Bacterial DNA Kit (Omega BIO-TEK) according to the manufacturer’s instructions. DNA yield and quality were measured and checked using a Qubit 3.0 Fluorometer (Invitrogen; Life Technologies) and gel electrophoresis.

Genome sequencing of all three bacterial genomes was conducted using the Illumina and PacBio platforms at Apical Scientific Sdn. Bhd (Malaysia) for *S. pseudonitzschiae* F5 and *Phaeobacter* sp. F10 and at Novogene (China) for *A. macleodii* F12. In brief, 100~1000 ng DNA was fragmented by acoustic disruption using a Covaris S220 system (Covaris, Woburn, MA) for Illumina sequencing. Subsequently, DNA libraries were built using a NEBNext® Ultra ™ II DNA Library Prep Kit (NEB E7645S/L, New England BioLabs® Inc.), and quantification and quality control of generated libraries were performed on an Agilent 2100 Bioanalyzer (Agilent Technologies, Santa Clara, CA). The libraries were then sequenced with TruSeq SBS Kit v4-HS reagents on an Illumina HiSeq 2500 platform (Illumina, San Diego, CA). For PacBio sequencing, DNA was sheared to ~15-20 kb using a Megarupter (Diagenode) and sequenced on a PacBio RS II system (Pacific Biosciences, Menlo Park, CA).

High-quality PacBio sub-reads were assembled by aligning the Illumina paired-end 150 bp reads with the Basic Local Alignment via Successive Refinement (BLASR) aligner, a tool that combines data from short read alignments with optimization methods from whole genome alignment (Chaisson and Tesler, 2012), and were trimmed using bbduk, aligned with bbmap, and the resulting sorted bam files and the Pacbio consensus reference were used as input to pilon (--genome genome.fasta, --bam input.bam) (Walker et al., 2014) for further error correction and polishing. The assemblies were further polished with arrow in Canu 1.7 with parameter corOutCoverage=60 (Koren et al., 2017). The largest circular contig in each assembly represented the chromosome size for each bacterium. In addition, plasmid contigs were BLASTed against closely related genomes to confirm these were indeed plasmids. For *S. pseudonitzschia* F5, we were not able to close the chromosome due to high number of repeats in the genome and thus exact chromosome size could not be determined (Table 1). The consensus reference was annotated by Prokka (Seemann, 2014) and RAST, and checked for completeness by BUSCO (Simão et al., 2015). Circa 1.2.1 (http://omgenomics.com/circa/) was used to draw chromosomes circos plot and chord diagram to compare the same gene locations in different genomes. All genome sequences and their annotations can be accessed at GenBank under the accession numbers WKFG01000000 and CP046140-CP046144.

Mining publicly available *Alteromonas* genomes for autoinducer synthase gene was performed on MicroScope (https://mage.genoscope.cns.fr/microscope/home/index.php). Identification and classification of transporter proteins in the three bacterial genomes was done using 6,097 membrane transport protein sequences downloaded from the Transporter Classification Database (TCDB) (Saier Jr et al., 2016). To this end, Gblast2 program (http://www.tcdb.org/labsoftware.php) with a cutoff e-value < 1e^−20^ was used and sequences with alignment scores <100 were removed. The chemotaxis, flagellar protein, pili, exopolysaccharides and quorum sensing related gene identities were confirmed by BLASTp with the model *Roseobacter* group strain *Ruegueria pomeroyi* DSS-3.

### QS genes analysis

The *luxR*-like and *luxI*-like genes of *S. pseudonitzschiae* F5 and *Phaeobacter* sp. F10 were predicted by RAST and Prokka. Maximum likelihood phylogenetic trees of multiple copies of LuxI-like, genetically linked LuxR-like and their internal transcribed spacer (ITS) sequences from *S. pseudonitzschiae* F5, *Phaeobacter* sp. F10, and 30 publicly accessible *Roseobacter* group genomes were constructed using PhyML 3.0 (Guindon et al., 2010) with HKY85 substitution model, which was selected according to SMS (Smart Model Selection in PhyML) (Lefort et al., 2017). These 30 strains were chosen based on several criteria: 1) Strains must have published information about at least two AHLs molecules, motility and attachment lifestyles, 2) All strains have whole genomes publicly available, and one of the following two criteria: 3) Strains are phylogenetically related to *S. pseudonitzschiae* F5 and *Phaeobacter* sp. F10, or 4) Strains are well studied as model roseobacters. The *luxRI* and ITS sequences of *Pseudomonas aeruginosa* PAO1 (RefSeq accession number NC_002516.2) were used as an outgroup. Phylogeny was tested with a fast likelihood-based method aBayes (Anisimova et al., 2011) with 100 bootstraps.

### AHLs extraction and identification

To extract AHL molecules, *S. pseudonitzschiae* F5 and *Phaeobacter* sp. F10 strains were re-plated from glycerol stocks and subsequently single picked colonies were inoculated overnight in 25-mL marine broth at 26℃ in the dark and shaking at 180 rpm. Overnight cultures were subsequently transferred to 1 L sterile marine broth in triplicates and grown under the same conditions for 20 hours. Finally, bacterial cells were removed by centrifugation (15 min, 7000 rpm) when the optical density (600 nm) reached 1.0. To further remove residual bacteria, the supernatant was further filtered through 0.2-μm polycarbonate membrane filters (Whatman, NJ, United States). Oasis HLB solid-phase extraction (SPE) cartridges (3 cc, 540 mg) were activated according to the manufacturer’s instructions and were subsequently used to remove organic molecules from the filtrates as previously described (Wang et al., 2017). Finally, extracts were eluted with 5 mL 0.1% (v/v) formic acid in methanol and dried using an evaporator (SpeedVac SC210A, Thermo Savant, Holbrook, NY, USA) and were stored at −20℃ for subsequent UHPLC-MS/MS analysis. AHL standards (Supplementary Table S4) were acquired from Cayman Chemicals and Sigma-Aldrich in order to help identify AHLs from *S. pseudonitzschiae* F5 and *Phaeobacter* sp. F10.

AHLs were analyzed using an Agilent 1290 HPLC system coupled to a Bruker Impact II Q-ToF-MS (Bruker, Germany). Metabolites were separated using a reversed-phase separation method. In RP mode, medium-polarity and non-polar metabolites were separated using an Eclipse Plus C_18_ column (50mm × 2.1mm ID) (Agilent, US). Chromatographic separation consisted of MilliQ-H_2_O + 0.2% formic acid (buffer A), and Acetonitrile + 0.2% formic acid (Buffer C) at a flowrate of 0.4 mL. The initial mobile phase composition was 90% A and 5% C followed by a gradient to 100% C over 18 min. The column was maintained at 100% C for 2 mins followed by cleaning with isopropanol for 3 min. The column was allowed to equilibrate for 4 min using the initial mobile phase composition. Detection was carried out in the positive ionization mode with the following parameters: Mass Range = 50 – 1300 m/z measured at 6 Hz, ESI source parameters: dry gas temperature = 220°C, dry gas flow = 10.0 l/min, Nebulizer pressure = 2.2 bar, Capillary V = 3000 V, end plate Offset: 500 V; MS-ToF tuning parameters: Funnel 1 RF = 150 Vpp, Funnel 2 RF = 200 Vpp, isCID Energy = 0 eV, Hexapole RF = 50 Vpp, Ion Energy = 4.0 eV, Low Mass = 90 m/z, Collision Energy = 7.0 eV, pre Pulse storage = 5 µs.

Data was processed and analyzed with Metaboscape 3.0 (Bruker, Bremen). Processing was conducted with the T-Rex3D algorithm with an intensity threshold of 500 and a minimum peak length of 10 spectra. Spectra were lock-mass calibrated, and features were only created if detected in a minimum of 3 samples. The presence of a specific AHL was determined by comparing the high-resolution parent ion mass, daughter ion masses and retention times of the purchased standards relative to bacterial extracts.

### Bacterial motility and biofilm formation assays

Bacterial motility assay was performed using semisolid (0.25% w/v) marine agar plates supplemented with a final concentration of 2 μM of each AHL or QS inhibitor (QSI) as described previously (Zan et al., 2012). The QSIs 2(5*H*)-furanone and (*Z*-)-4-Bromo-5-(bromomethylene)-2(5*H*)-furanone (furanone C-30) were purchased from Sigma-Aldrich. *S. pseudonitzschiae* F5, *Phaeobacter* sp. F10, and *A. macleodii* F12 strains were grown in marine broth overnight, and then were gently inoculated using a sterilized toothpick into the center of the agar surface. Triplicate plates were incubated at 26°C for 3 days after which motility plates were observed using the Uvitec Cambridge Fire-reader imaging system. The proportion of motile area (percent motility) was measured using ImageJ software (http://rsb.info.nih.gov/ij/). Bacterial motility is characterized by a circular swimming zone in the motility assay or by dendritic morphology, which is typical for swarming motility caused by a locally restricted movement on agar plates (Michael et al., 2016).

Bacterial biofilm formation was quantified for *S. pseudonitzschiae* F5, *Phaeobacter* sp. F10, and *A. macleodii* F12 strains using the crystal violet assay (O’toole and Kolter, 1998). Briefly, bacteria were cultured in marine broth overnight and then diluted to ~1×10^5^ cells/mL. 100 μL aliquots were transferred to a 96-well suspension culture plate (Greiner bio-one, CELLSTAR, Monroe, NC, USA). 10 μg/mL AHL or QSI were added to each well respectively in triplicates as final concentration, which is optimized from previous studies (Ren et al., 2001; Zhu et al., 2019) while controls received no AHL or QSI addition. The plate was incubated at 26℃ for 24 h and subsequently 25 μL of 0.1% crystal violet was added to each well and incubated at room temperature for 15 min. Wells were rinsed twice with 150 μL Milli-Q water to remove free-living bacteria and then dried at 60℃ for 10 min. Finally, stained biomass was solubilized in 95% ethanol for 1 h and absorbance readings were measured using a plate reader (BioTek, Winooski, VT) at 600 nm. Unpaired t-test was used to compare the significant differences of control and other variables in bacterial swarming and biofilm formation.

## Acknowledgements

We would like to thank the China Scolarship Council for the financial support of C.F. This research was partially carried out using the Core Technology Platforms resources at New York University Abu Dhabi. This work was funded by an NYU Abu Dhabi grant (AD179) to S.A.A.

