## Supplementary information for "Quorum Sensing Regulates ‘swim-or-stick’ Lifestyle in the Phycosphere"

Supplementary Figures S1-S5

Supplementary Tables S1-S4

References

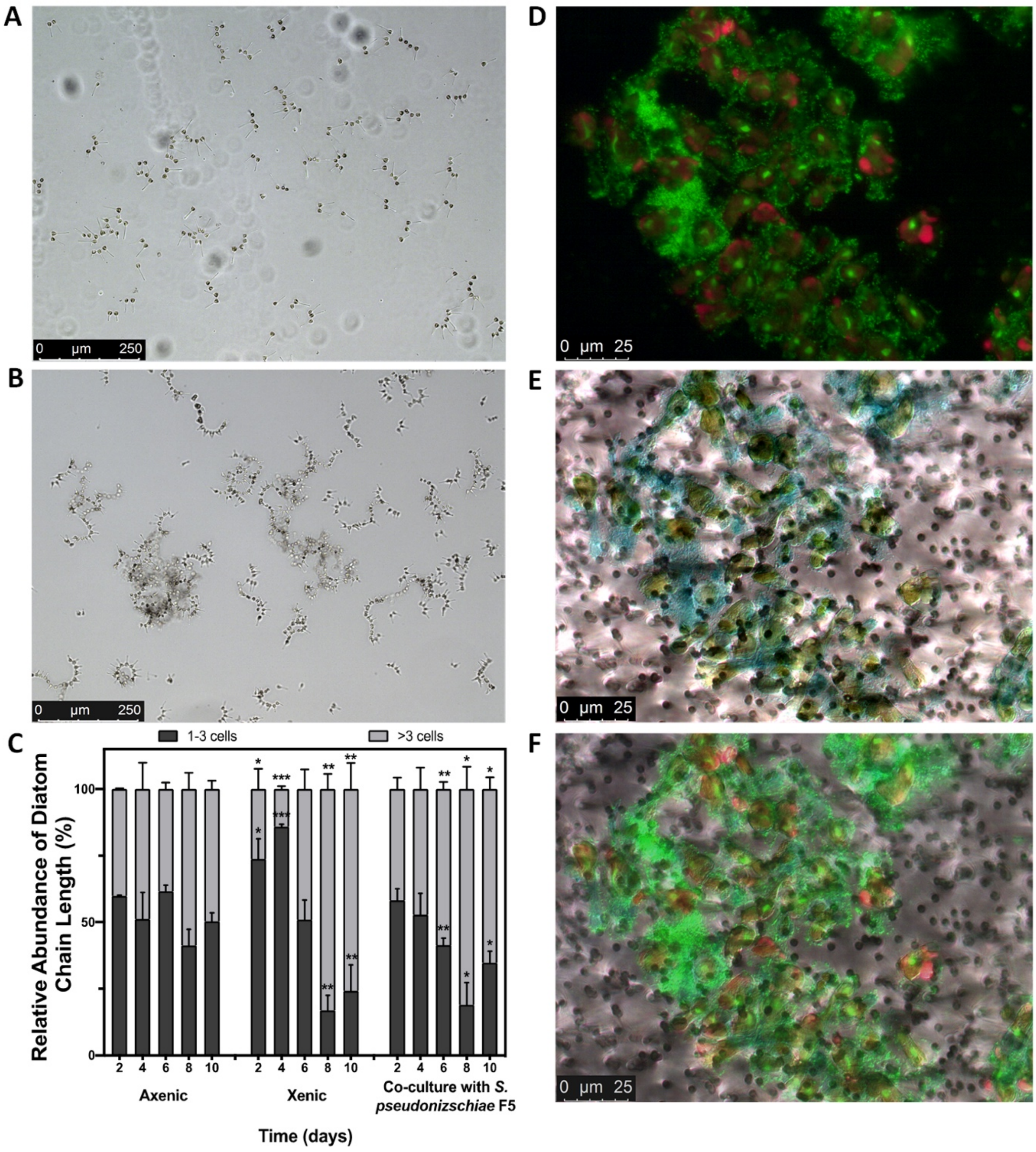


**Supplementary Fig. S1.** Bacteria influence chain length and display strong attachment to *Asterionellopsis glacialis* A3. Bright-field microscopic images of axenic **(A)** and xenic **(B)** *A. glacialis* A3 at equivalent cell densities (~$1.5\times10^5$ cells/mL) displaying a general trend towards longer chains in xenic cultures. **(C)** The relative abundance of different diatom chain lengths in cultures of axenic, xenic *A. glacialis* A3 and co-cultures of *A. glacialis* and *S. pseudonitzschiae* F5. Error bars represent standard deviation (SD) of triplicate cultures. Statistical significance is denoted by * $p<0.05$, ** $p<0.01$, and *** $p<0.001$ for xenic and coculture samples compared to axenic cultures at each time point. **(D-F)** Micrographs of xenic *A. glacialis* A3 cells and attached bacteria: **(D)** epifluorescence field, **(E)** bright field, and **(F)** merged image of (D) and (E). Red color corresponds to diatom chlorophyll *a*

autofluorescence, while green color indicates nucleic acids of diatoms and bacteria. Blue color in (E) and (F) corresponds to diatom transparent exopolymeric particles (TEP) stained with alcian blue.

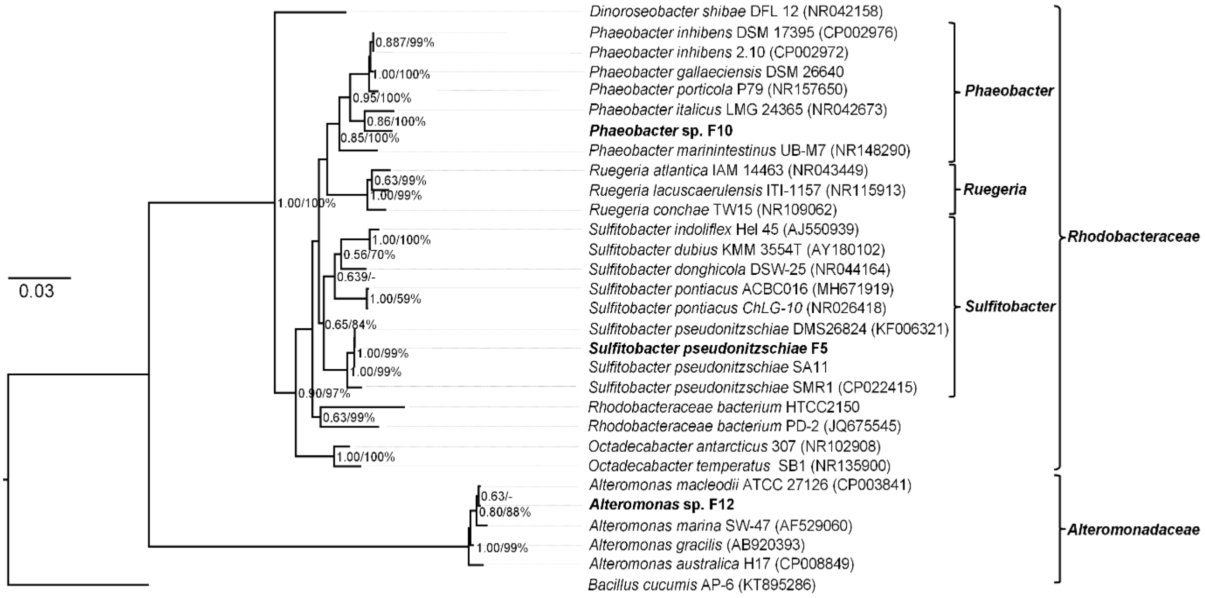


**Supplementary Fig. S2.** Bayesian phylogenetic tree of the 16S rRNA gene of cultivable bacteria from the *A. glacialis* A3 microbial consortium. Three bacteria isolated from *A. glacialis* A3 are highlighted in bold. Genera and families of the different bacteria are indicated to the right. Posterior probabilities and ML bootstrap values are listed adjacent to branches, respectively. GenBank accession numbers are denoted in parentheses.

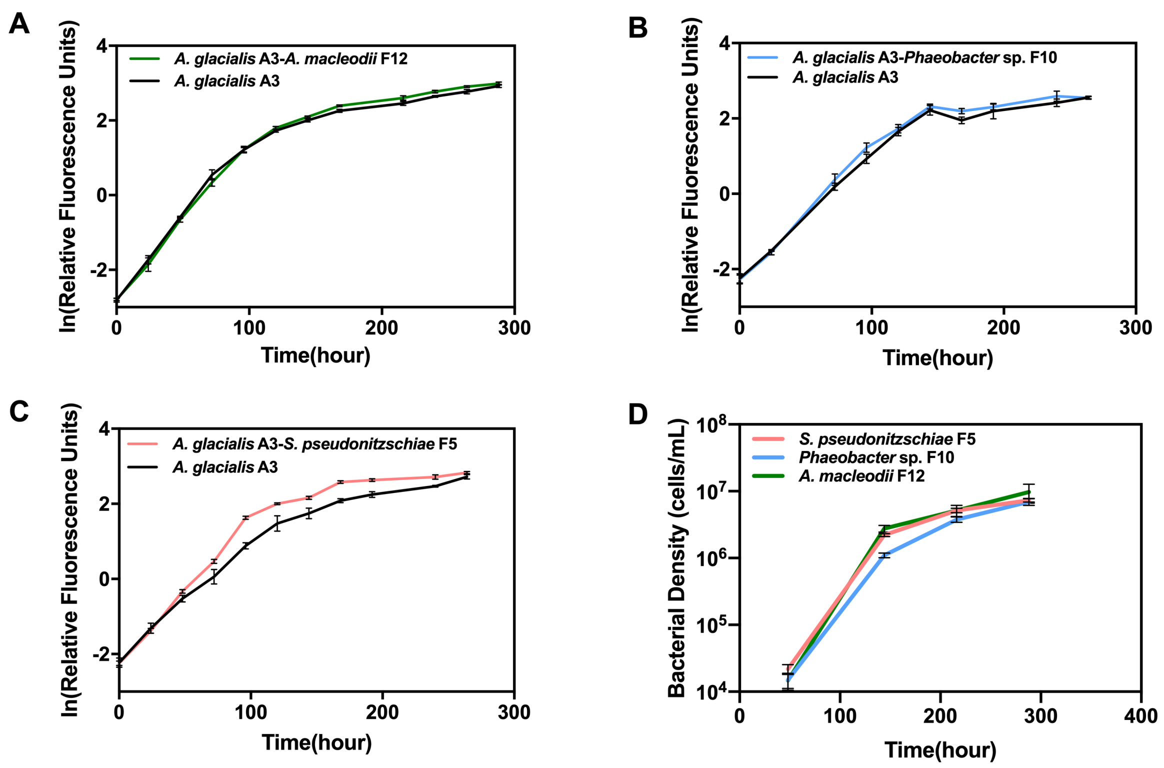


**Supplementary Fig. S3.** Effects of bacterial isolates on the specific growth of *A. glacialis* A3. Co-culture of *A. glacialis* A3 with **(A)** *Alteromonas macleodii* F12, **(B)** *Phaeobacter* sp. F10**,** and **(C)** *S. pseudonitzschiae* F5**. (D)** Cell density of each bacterium co-cultured with A3. Error bars represent S.D. of six replicates.

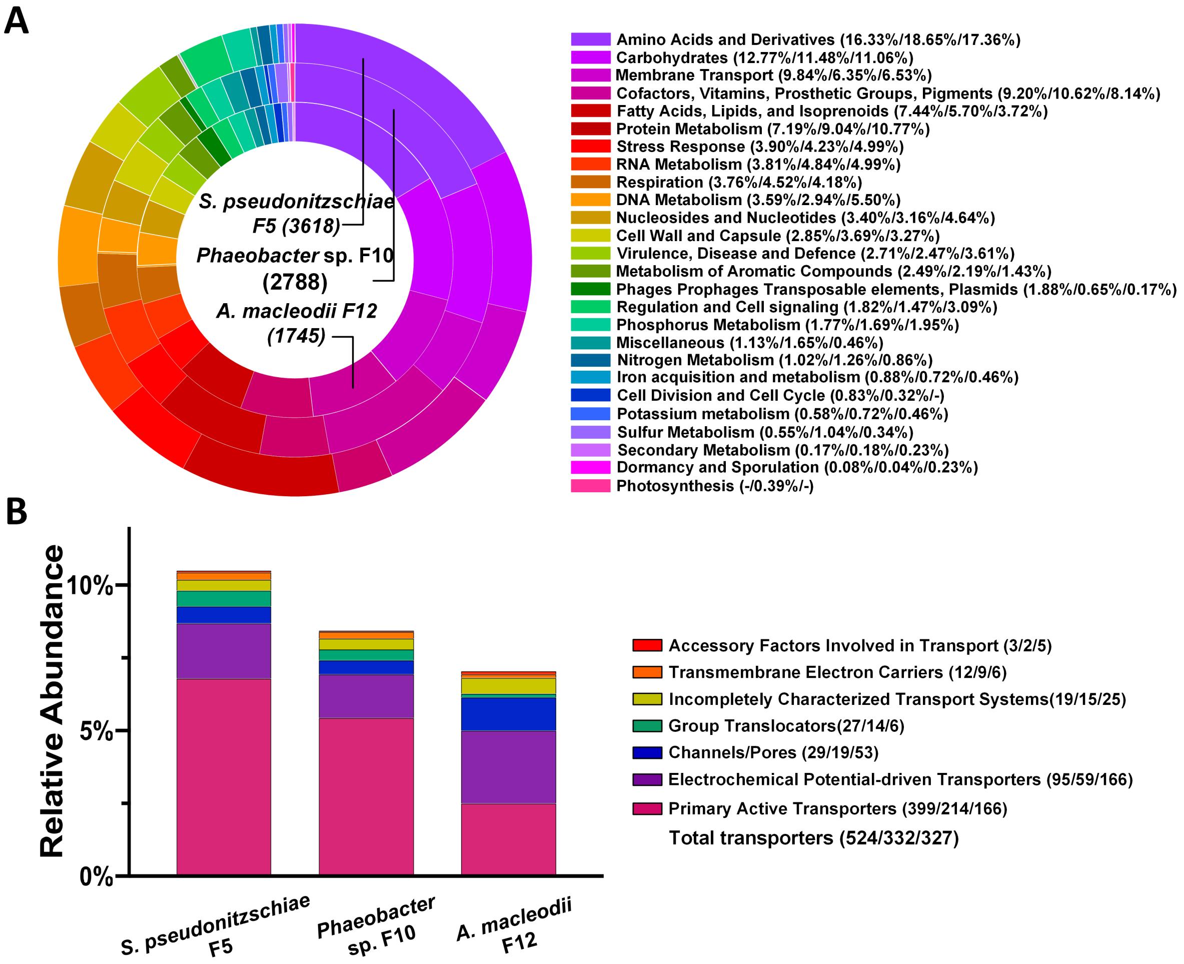


**Supplementary Fig. S4.** Genome-wide analysis of annotated genes for *S. pseudonitzschiae* F5, *Phaeobacter* sp. F10, and *A. macleodii* F12. **(A)** Number of total annotated genes for each bacterium is listed in parentheses on the pie chart. Proportion of genes in each category belonging to *S. pseudonitzschiae* F5 (left), *Phaeobacter* sp. F10 (middle) and *A. macleodii* F12 (right) are listed next to each category. **(B)** Normalized number of total putative transporters to the genome size of each bacterium. Number of genes in each category belonging to *S. pseudonitzschiae* F5 (left), *Phaeobacter* sp. F10 (middle) and *A. macleodii* F12 (right) are listed next to each category.

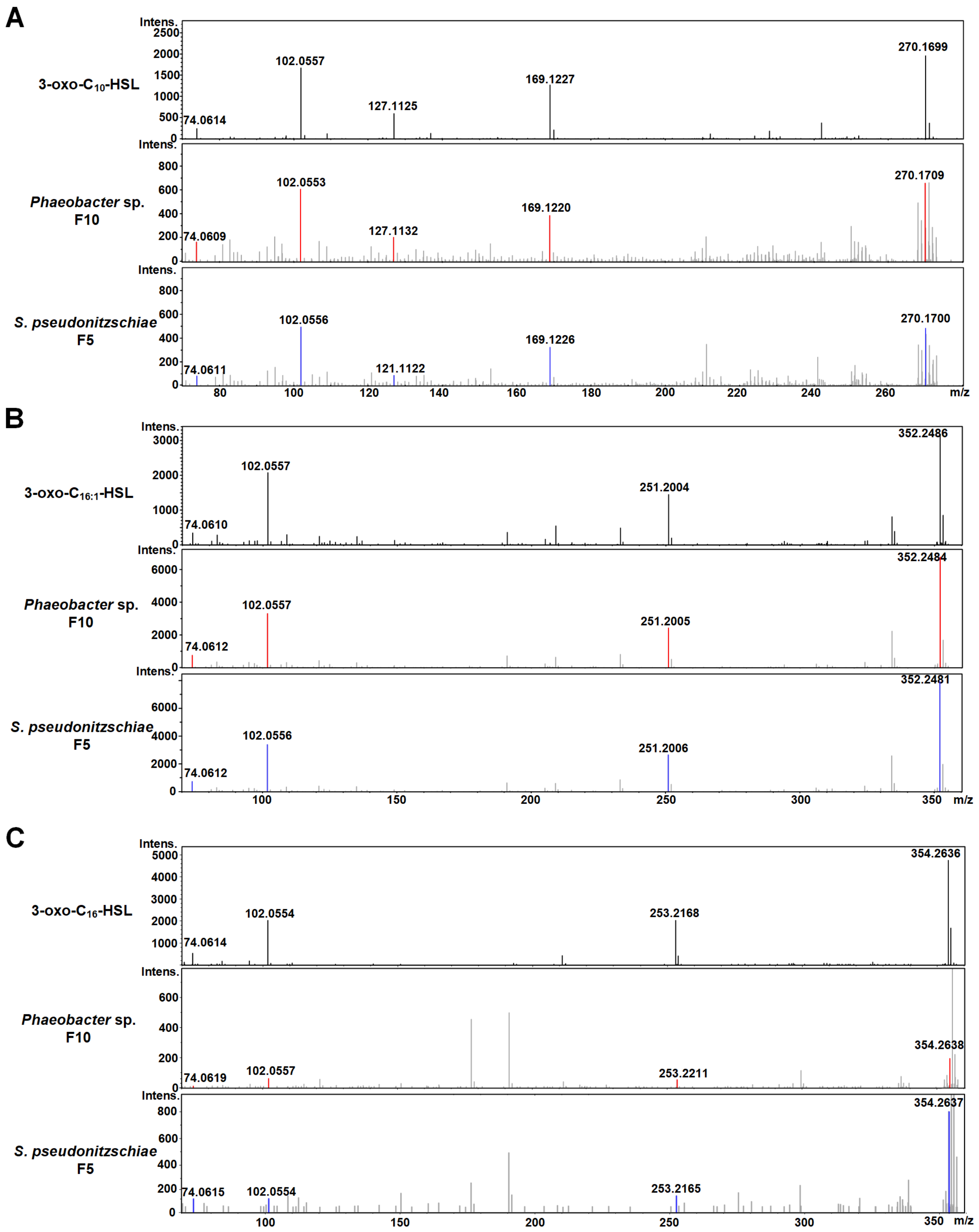


**Supplementary Fig. S5.** UHPLC-MS/MS spectra of three AHLs from purchased standards, supernatants of *S. pseudonitzschiae* F5 or *Phaeobacter* sp. F10. MS/MS spectra of **(A)** 3-oxo-$C_{10}$-HSL, **(B)** 3-oxo-$C_{16:1}$-HSL and **(C)** 3-oxo-$C_{16}$-HSL are shown and specific fragment ion peaks are labeled with their m/z values in standards (black), *S. pseudonitzschiae* F5 (red), and *Phaeobacter* sp. F10 (blue).

**Supplementary Table S1.** Putative IAA biosynthesis pathway in *S. pseudonitzschiae* F5

| Gene annotation | Feature ID | % ID* | Bit score* |
|---|---|---|---|
| IAM hydrolase [EC:3.5.1.4] | fig\|1402135.21.peg.3560 | 97.4 | 523 |
| Tryptophan 2-monooxygenase | fig\|1402135.21.peg.3559 | 100 | 1148 |
| Putative IpyA decarboxylase | fig\|1402135.21.peg.2326 | 98.9 | 946 |
| Aldehyde dehydrogenase (NAD+) | fig\|1402135.21.peg.2628 | 99.8 | 981 |
| Nitrile hydratase subunit alpha | fig\|1402135.21.peg.363 | 99.5 | 375 |
| Nitrile hydratase subunit beta | fig\|1402135.21.peg.362 | 100 | 452 |

*Identity (% ID) and bit score refer to identity of homologs in *S. pseudonitzschia* F5 compared to *S. pseudonitzschiae* SA11.

**Supplementary Table S2.** Homolog genes of *S. pseudonitzschiae* F5, *Phaeobacter* sp. F10, and *A. macleodii* F12 depicted in Figure 2 and their similarity to homologs in the genome of the model bacterium *Ruegeria pomeroyi* DSS-3. Complete annotation of each genome is provided as a separate supplementary file.

| Gene | Annotation | *S. pseudonitzschiae* F5 | % ID | Bit score | *Phaeobacter* sp. F10 | % ID | Bit score | *A. macleodii* F12 | % ID | Bit score |
|---|---|---|---|---|---|---|---|---|---|---|
| **Chemotaxis** | | | | | | | | | | |
| *cheR* | Chemotaxis protein methyltransferase | fig\|1402135.21.peg.3438 | -[§] | - | fig\|6666666.382773.peg.987<br>fig\|6666666.382773.peg.2269 | - | - | fig\|28108.372.peg.1943 | - | - |
| | Chemotaxis protein histidine kinase and related kinases | fig\|1402135.21.peg.1719 | 53.4 | 111 | fig\|6666666.382773.peg.1026 | 65.7 | 134 | - | | - |
| *cheB* | Chemotaxis response regulator protein-glutamate methylesterase CheB | - | | - | fig\|6666666.382773.peg.981<br>fig\|6666666.382773.peg.2270 | -<br>33.3 | -<br>55.8 | fig\|28108.372.peg.361<br>fig\|28108.372.peg.1933 | -<br>34.9 | -<br>63.2 |
| *cheY* | Chemotaxis regulator - transmits chemoreceptor signals to flagelllar motor components CheY | fig\|1402135.21.peg.3437 | - | - | fig\|6666666.382773.peg.988 | | - | fig\|28108.372.peg.1936<br>fig\|28108.372.peg.1704 | | - |
| *cheW* | Positive regulator of CheA protein activity | fig\|1402135.21.peg.3439 | | - | fig\|6666666.382773.peg.986<br>fig\|6666666.382773.peg.2266<br>fig\|6666666.382773.peg.2268 | | - | fig\|28108.372.peg.1930 | | - |
| *cheA* | Signal transduction histidine kinase CheA | fig\|1402135.21.peg.3440 | | - | fig\|6666666.382773.peg.985<br>fig\|6666666.382773.peg.2265 | | -<br>- | fig\|28108.372.peg.1934 | | - |
| *cheD* | Chemotaxis protein CheD | fig\|1402135.21.peg.3435 | | - | fig\|6666666.382773.peg.980 | | - | - | | - |
| **Flagellar Proteins** | | | | | | | | | | |
| *motB* | Flagellar motor rotation protein MotB | fig\|1402135.21.peg.769<br>fig\|1402135.21.peg.1597 | 55.8<br>53.8 | 626<br>223 | fig\|6666666.382773.peg.243<br>fig\|6666666.382773.peg.909 | 66.5<br>60.7 | 750<br>232 | fig\|28108.372.peg.1443 | - | - |

| | | | | | | | | | | |
|---|---|---|---|---|---|---|---|---|---|---|
| | MotA/TolQ/ExbB proton channel family protein, probably associated with flagella | fig\|1402135.21.peg.770 | 69.5 (hp) ¶ | 515 | fig\|6666666.382773.peg.244 | 75.7 (hp) | 569 | - | | - |
| *flgN* | Flagellar biosynthesis protein FlgN | fig\|1402135.21.peg.1599 | 46.2 (hp) | 108 | fig\|6666666.382773.peg.902 | 46.2 (hp) | 105 | - | | - |
| *fliG* | Flagellar motor switch protein FliG | fig\|1402135.21.peg.134 | 56.8 | 328 | fig\|6666666.382773.peg.2790 | 60.7 | 384 | - | | |
| *flgE* | Flagellar hook protein FlgE | fig\|1402135.21.peg.1598 | 65.4 | 566 | - | | | fig\|28108.372.peg.1939 | - | - |
| *flgJ* | Flagellar protein FlgJ, putative | fig\|1402135.21.peg.1600 | 65.7 | 95.5 | fig\|6666666.382773.peg.901 | 73.5 | 103 | - | | - |
| | Flagellar motor switch protein | fig\|1402135.21.peg.210 | 30.8 (hp) | 117 | fig\|6666666.382773.peg.2711 | 36.5 (hp) | 159 | - | | - |
| *flgD* | Flagellar basal-body rod modification protein FlgD | fig\|1402135.21.peg.1602 | 56.9 | 229 | fig\|6666666.382773.peg.899 | 51.5 | 213 | - | | - |
| **Pili** | | | | | | | | | | |
| | bacterial type II/III secretion system protein | fig\|1402135.21.peg.2422 | 33.1 | 185 | fig\|6666666.382773.peg.2057 | 75.4 | 444 | - | | - |
| | Predicted ATPase with chaperone activity, associated with Flp pilus assembly | fig\|1402135.21.peg.2680 | 82.3 (hp) | 758 | fig\|6666666.382773.peg.2048 | 84.8 (hp) | 786 | - | | - |
| *tadC* | type II/IV secretion system protein, TadC subfamily | fig\|1402135.21.peg.2426<br>fig\|1402135.21.peg.2685<br>fig\|1402135.21.peg.2684 | -<br>78.1<br>68.9 | -<br>514<br>436 | fig\|6666666.382773.peg.2053<br>fig\|6666666.382773.peg.2052 | 75.2<br>72.4 | 504<br>484 | - | | - |
| *tadA* | type II/IV secretion system protein, TadA subfamily | fig\|1402135.21.peg.2425<br>fig\|1402135.21.peg.2686 | 50.5<br>82.5 | 391<br>784 | fig\|6666666.382773.peg.2054 | 86.0 | 827 | - | | - |
| | Type II/IV secretion system ATPase | fig\|1402135.21.peg.2687<br>fig\|1402135.21.peg.2424 | 72.5<br>- | 632<br>- | fig\|6666666.382773.peg.2055 | 74.0 | 649 | - | | - |

| | | | | | | | | | | |
|---|---|---|---|---|---|---|---|---|---|---|
| | TadZ/CpaE, associated with Flp pilus assembly | | | | | | | | | |
| *tadG* | Flp pilus assembly protein TadG | fig\|1402135.21.peg.893 | 39.1 | 132 (hp) | fig\|6666666.382773.peg.2304 | 44.6 | 171 (hp) | - | | - |
| | ATP-dependent helicase, DEAD/DEAH box family, associated with Flp pilus assembly | fig\|1402135.21.peg.2679 | 81.8 | 1344 | fig\|6666666.382773.peg.2047 | 77.4 | 1279 | - | | - |
| *tadD* | Flp pilus assembly protein TadD, contains TPR repeat | fig\|1402135.21.peg.2682<br>fig\|1402135.21.peg.2683 | 75.8<br>59.1 | 449<br>218 | fig\|6666666.382773.peg.2050;<br>fig\|6666666.382773.peg.2051 | 71.7<br>58.2 | 418<br>210 | - | | - |
| | Flp pilus assembly protein RcpC/CpaB | fig\|1402135.21.peg.2690 | 68.4 | 395 | fig\|6666666.382773.peg.2058 | 70.9 | 416 | - | | - |
| | Flp pilus assembly protein, pilin Flp | fig\|1402135.21.peg.2420<br>fig\|1402135.21.peg.2692 | -<br>- | -<br>- | fig\|6666666.382773.peg.2060 | - | - | fig\|28108.372.peg.1117 | | - |
| | Type IV prepilin peptidase TadV/CpaA | fig\|1402135.21.peg.2681 | 51.9 | 154 | fig\|6666666.382773.peg.2049 | 48.6 | 123 | - | | - |
| **Exopolysaccharides** | | | | | | | | | | |
| | bacterial sugar transferase | fig\|1402135.21.peg.2078<br>fig\|1402135.21.peg.1856 | 48.1<br>35.8 | 253<br>224 | fig\|6666666.382773.peg.3316 | 65.6 | 253 | fig\|28108.372.peg.1531 | 34.9 | 244 |
| | polysaccharide biosynthesis/export protein | fig\|1402135.21.peg.873 | 71.9 | 377 | fig\|6666666.382773.peg.2324 | 82.4 | 375 | - | | - |
| *kpsC* | capsular polysaccharide export protein KpsC | fig\|1402135.21.peg.874 | 67.0 | 667 | fig\|6666666.382773.peg.2323 | 70.4 | 667 | - | | - |
| | Glycosyl transferase, group 2 family protein | fig\|1402135.21.peg.303 | 54.6 | 628 | fig\|6666666.382773.peg.420 | 50.1 | 637 | - | | - |
| *kpsS* | Capsular polysaccharide export system protein KpsS | fig\|1402135.21.peg.872 | 75.1 | 430 | fig\|6666666.382773.peg.2325 | 77.7 | 430 | - | | - |

| | | | | | | | | | | |
|---|---|---|---|---|---|---|---|---|---|---|
| | Glycosyl transferase, group 1 family protein | fig\|1402135.21.peg.2297 | 72.6 | 413 | fig\|6666666.382773.peg.732 | 74.3 | 413 | - | | - |
| **Quorum sensing** | | | | | | | | | | |
| *luxRI* cassette (B group) | Autoinducer-binding transcriptional regulator and adjacent autoinducer synthesis | fig\|1402135.21.peg.2496<br>fig\|1402135.21.peg.2495 | 67.9<br>68.1 | 304<br>344 | fig\|6666666.382773.peg.1880<br>fig\|6666666.382773.peg.1879 | 73.6<br>73.2 | 318<br>381 | - | | - |
| *luxRI* cassette | Autoinducer-binding transcriptional regulator and adjacent autoinducer synthesis | fig\|1402135.21.peg.2198<br>fig\|1402135.21.peg.2199 | 71.2<br>60.8 | 313<br>290 | - | - | - | | | |
| Orphan *luxR* | Autoinducer-binding transcriptional regulator, LuxR family | fig\|1402135.21.peg.3467 | 71.5 | 382; | fig\|6666666.382773.peg.2975 | 68.3 | 363 | - | | - |
| | DNA-binding response regulator, LuxR family | fig\|1402135.21.peg.2410<br>fig\|1402135.21.peg.3268 | 52.2<br>76.6 | 204<br>325 | fig\|6666666.382773.peg.3362<br>fig\|6666666.382773.peg.2638 | 62.6<br>83.2 | 249<br>353 | - | | - |
| | Transcriptional regulator, LuxR family | - | | - | fig\|6666666.382773.peg.668<br>fig\|6666666.382773.peg.2699 | 44.3<br>38.0 | 108<br>108 | fig\|28108.372.peg.736 | 31.7 | 69.3 |

Further confirmation for gene annotations were acquired by blasting genes against the model *Roseobacter* group bacterium *Ruegeria pomeroyi* DSS-3 (Hit score > 50)

§ Indicates no hits were found in the *R. pomeroyi* DSS3 genome.

¶ Indicates that the gene annotation in *R. pomeroyi* DSS3 is 'hypothetical protein'.

**Supplementary Table S3.** AHL molecules listed in Figure 5.

| Strains | No. of *luxI*s | AHLs | References |
|---|---|---|---|
| *Phaeobacter* sp. F10 | 1 | 3-oxo-$C_{10}$-HSL, 3-oxo-$C_{16}$-HSL, 3-oxo-$C_{16:1}$-HSL | This study |
| *Rhodobacterales bacterium* Y4I | 2 | $C_8$-HSL, 3-OH-$C_{12:1}$-HSL | [1] |
| *Phaeobacter gallaeciensis* DSM 26640 | 3 | 3-OH-$C_{10}$-HSL, $C_{12:1}$-HSL, $C_{14:2}$-HSL, $C_{16:1}$-HSL, $C_{16:2}$-HSL, 3-oxo-$C_{16}$-HSL, $C_{18:1}$-HSL, $C_{18:2}$-HSL, | [4] |
| *Phaeobacter gallaeciensis* DSM 17395 | 2 | 3-OH-$C_{10}$-HSL, $C_{16}$-HSL, $C_{16:1}$-HSL, $C_{18:1}$-HSL | [5] |
| *Phaeobacter inhibens* 2.10 | 3 | 3-OH-$C_{10}$-HSL, 3-oxo-$C_{10}$-HSL, $C_{12:2}$-HSL, $C_{16}$-HSL, $C_{16:1}$-HSL, $C_{18:1}$-HSL | [5] |
| *Ruegeria* sp. KLH11 | 3 | 3-OH-$C_{14}$-HSL, 3-OH-$C_{14:1}$-HSL, 3-OH-$C_{12}$-HSL | [9] |
| *Dinoroseobacter shibae* DFL 12 | 3 | $C_{14:1}$-HSL, 3-oxo-$C_{14}$-HSL, $C_{18:1}$-HSL, $C_{18:2}$-HSL | [5] |
| *Rhodobacteraceae bacterium* PD-2 | 2 | 3-oxo-$C_8$-HSL, 3-oxo-$C_{10}$-HSL | [14] |
| *Sulfitobacter pseudonitzschiae* F5 | 2 | 3-oxo-$C_{10}$-HSL, 3-oxo-$C_{16}$-HSL, 3-oxo-$C_{16:1}$-HSL | This study |
| *Pseudomonas aeruginosa* PAO1 | 2 | $C_4$-HSL, 3-oxo-$C_{12}$-HSL | [13] |

**Supplementary Table S4.** Purchased AHLs standards used to identify AHLs produced by the roseobacters strains using HPLC-MS/MS.

| AHL standards | Future ID |
|---|---|
| Acetyl-L-Homoserine lactone | $C_2$-HSL |
| N-butyryl-L-Homoserine lactone | $C_4$-HSL |
| N-hexanoyl-L-Homoserine lactone | $C_6$-HSL |
| N-(β-Ketocaproyl)-DL-homoserine lactone | 3-oxo-$C_6$-HSL |
| N-phenylacetyl-L-Homoserine lactone | $C_8$-HSL |
| N-(3-Oxooctanoyl)-L-homoserine lactone | 3-oxo-$C_8$-HSL |
| N-(3-Oxodecanoyl)-L-homoserine lactone | 3-oxo-$C_{10}$-HSL |
| N-dodecanoyl-L-Homoserine lactone | $C_{12}$-HSL |
| N-(3-Oxododecanoyl)-L-homoserine lactone | 3-oxo-$C_{12}$-HSL |
| N-tetradecanoyl-L-Homoserine lactone | $C_{14}$-HSL |
| N-3-oxo-tetradecanoyl-L-Homoserine lactone | 3-oxo-$C_{14}$-HSL |
| N-hexadecanoyl-L-Homoserine lactone | $C_{16}$-HSL |
| N-3-oxo-hexadec-11(Z)-enoyl-L-Homoserine lactone | 3-oxo-$C_{16:1}$-HSL |
| N-3-oxo-hexadecanoyl-L-Homoserine lactone | 3-oxo-$C_{16}$-HSL |
| N-octadecanoyl-L-Homoserine lactone | $C_{18}$-HSL |

**Reference (including full references in Figure 5 and Supplementary Table S3)**
